# Metabolic control of histone acetylation for precise and timely regulation of minor ZGA in early mammalian embryos

**DOI:** 10.1101/2022.06.10.495405

**Authors:** Jingyu Li, Jiaming Zhang, Weibo Hou, Xu Yang, Xiaoyu Liu, Yan Zhang, Meiling Gao, Ming Zong, Zhixiong Dong, Zhonghua Liu, Jingling Shen, Weitao Cong, Chunming Ding, Shaorong Gao, Guoning Huang, Qingran Kong

## Abstract

Metabolism feeds into the regulation of epigenetics via metabolic enzymes and metabolites. However, metabolic features, and their impact on epigenetic remodeling during mammalian pre-implantation development, remain poorly understood. In this study, we established the metabolic landscape of mouse pre-implantation embryos from zygote to blastocyst, and quantified some absolute carbohydrate metabolites. We integrated these data with transcriptomic and proteomic data, and discovered the metabolic characteristics of the development process, including the activation of methionine cycle from 8-cell embryo to blastocyst, high glutaminolysis metabolism at blastocyst stage, enhanced TCA cycle activity from the 8-cell embryo stage, and active glycolysis in the blastocyst. We further demonstrated that oxidized nicotinamide adenine dinucleotide (NAD^+^) synthesis is indispensable for mouse pre-implantation development. Mechanistically, in part, NAD^+^ is required for the exit of minor zygotic gene activation (ZGA) by cooperating with SIRT1 to remove zygotic H3K27ac. In human, NAD^+^ supplement can promote the removal of zygotic H3K27ac and benefit pre-implantation development. Our findings demonstrate that precise and timely regulation of minor ZGA is controlled by metabolic dynamics, and enhance our understanding of the metabolism of mammalian early embryos.

**Highlights:**

- We identified the metabolic features of mouse pre-implantation embryo development.
- NAD^+^ synthesis is indispensable for mouse embryo pre-implantation development.
- NAD^+^-mediated erasure of zygotic H3K27ac is required for the exit of minor ZGA.
- NAD^+^ supplement promotes the removal of zygotic H3K27ac and benefits preimplantation development of human embryos.

## Introduction

Mammalian life begins with the fertilization of an oocyte by a sperm; during this process, these two terminally differentiated germ cells are converted into a totipotent zygote.^1^ The zygote undergoes pre-implantation development, giving rise to the blastocyst. This process involves a series of significant biological events, including zygotic gene activation (ZGA) followed by the first cell-fate decision and lineage-specific differentiation.^2,3^ In mice, the initial ZGA occurs between the S phase of the zygote and G1 of the early 2-cell embryo, and is designated as the minor ZGA to discriminate it from the burst of transcription that occurs during the mid-to-late 2-cell stage, which is designated as the major ZGA.^4,5^ Minor ZGA precedes major ZGA, and timely regulation of minor ZGA is crucial for pre-implantation development;^6^ this requires a functional and precise regulatory network.

Cellular metabolism is the foundation of all biological activity.^7^ Changes in metabolism are directly linked to changes in chromatin and DNA state.^8-10^ The importance of metabolism for controlling pre-implantation embryo development was established several decades ago through identification of the conditions that allow early embryos to grow outside of the oviduct.^11-13^ Recent research has shown that pyruvic acid is essential for initiating ZGA and selective translocation of key mitochondrial tricarboxylic acid (TCA) cycle proteins to the nucleus, indicating that this process allows epigenetic remodeling during ZGA.^14^ In morula, a glucose-mediated signaling process has been demonstrated to be critical for controlling expression of the transcription factors necessary for trophectoderm (TE) cell differentiation.^15^ Many studies have also suggested that balanced and timely metabolism is essential for pre-implantation embryo development competence.

Although intrinsic control of embryogenesis by intracellular metabolites and metabolic pathways has been studied extensively, previous studies have identified only some of the cellular pathways involved in this process.^16^ The establishment of global metabolic patterns could lead to a more complete understanding of pre-implantation embryo development. Therefore, in the study, we established a dynamic metabolome profile of mouse pre-implantation embryos from the zygote to blastocyst stages, and bolstered the metabolomics studies with quantitative transcriptomic and proteomic analysis. We also demonstrated that the combination of the prominent metabolic cofactor nicotinamide adenine dinucleotide (NAD^+^) and deacylase SIRT1 can remove zygotic H3K27ac, for precise and timely regulation of minor ZGA; this is essential for mouse pre-implantation embryo development. The results of this study will provide a reference for further studies of metabolism in early mammalian embryos, and new opportunities for the discovery of biomarkers to predict and improve pre-implantation embryo quality.

## Results

### Establishing temporal metabolome profiles of mouse pre-implantation embryo development

To systematically monitor metabolomic profiles during mouse pre-implantation embryo development, we isolated a large number of mouse embryos (36,000) from the zygote to blastocyst stages, and performed untargeted metabolomic analysis of the intracellular metabolome using ultra-high-performance liquid chromatography coupled with hybrid quadrupole time-of-flight mass spectrometry (UHPLC-Q-TOF-MS), each with six biological replicates (Fig. 1a). A total of 246 metabolites were identified (Supplementary Table S1). To identify the metabolites specific to each stage, we intersected metabolites with levels higher than the overall average of metabolites from all samples, and found that 4, 4, 6, 8, 7, and 14 metabolites were abundant at the zygote, 2-cell, 4-cell, 8-cell, morula, and blastocyst stages, respectively (Fig. 1b). Next, the metabolites in each stage were classified into three groups according to metabolite level: high, intermediate, or low. Some high metabolites decreased from the zygote to 2-cell embryo stages, suggesting the consumption of maternal metabolites accumulated during oogenesis. Across the continuous stages, metabolites had mainly low to high levels. After 4-cell embryo stage, a large number of low metabolites were observed in zygote and 2-cell embryo, trending toward high metabolites, especially at the morula stage and further enhanced in blastocyst (Fig. 1c), demonstrating metabolic activities were remarkably increased at the later stages. Uniform manifold approximation and projection (UMAP) was applied to the metabolites to distinguish the embryos according to their developmental stages, illustrating the metabolic dynamics during mouse pre-implantation embryo development. (Fig. 1d). This analysis revealed the highly dynamic changes in the metabolome during pre-implantation development from the earlier to later stages.

**Fig. 1.**
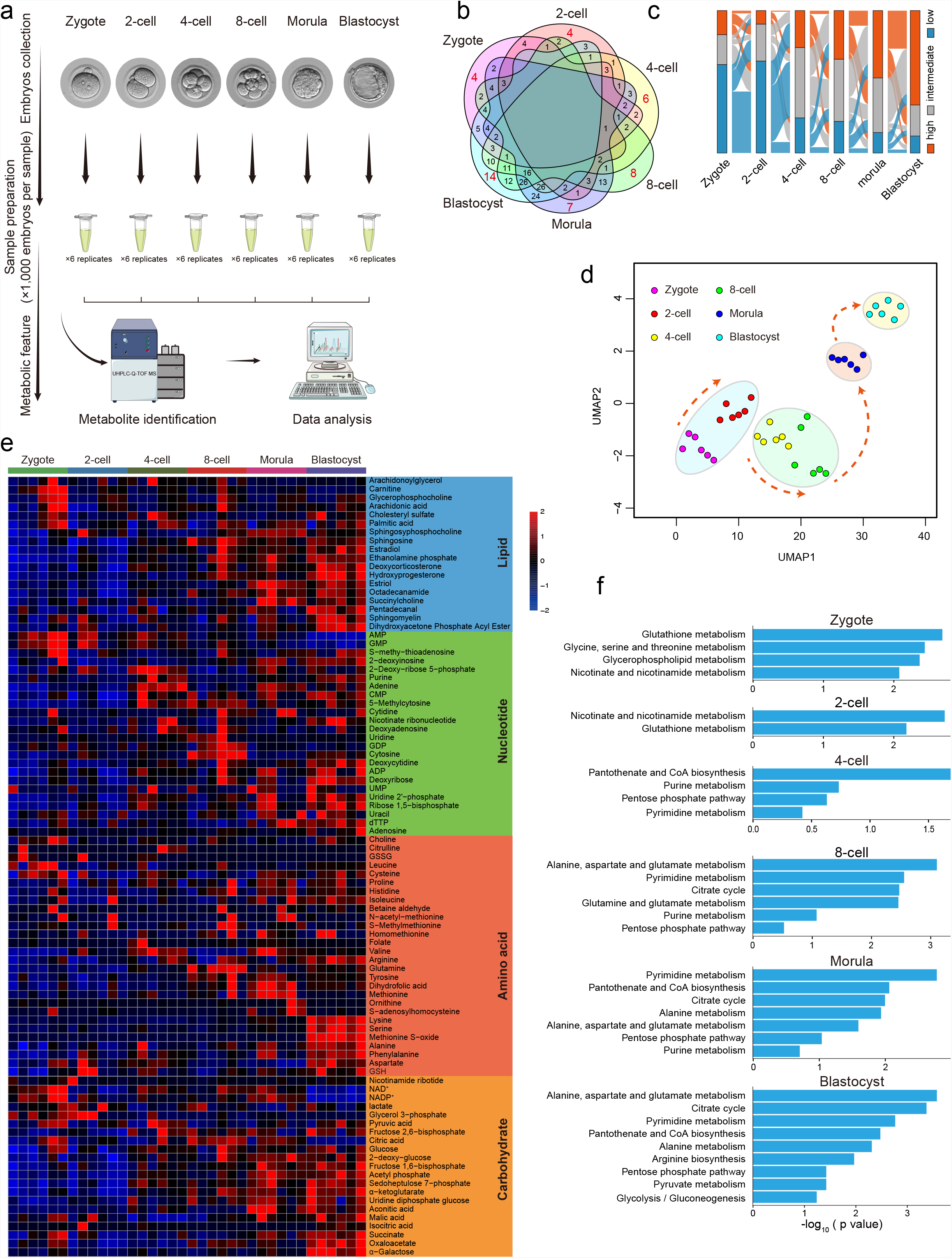
Metabolomic profiling of mouse pre-implantation embryo development. **a** Schematic overview of mouse embryo preparation and metabolome profiling during mouse pre-implantation embryo development. **b** Venn diagram showing the overlap among metabolites across all stages. The number of significantly abundant metabolites is shown in red. **c** Alluvial plots of the global dynamics of high, intermediate, and low metabolites based on their levels during early embryo development. **d** UMAP analysis. Metabolites marked with the same color represent six replicates of the same stage. **e** Heatmap of metabolite dynamics during mouse embryo development, classified according to metabolic pathways. **f** KEGG analysis of abundant metabolites in each stage.

### Metabolic features during mouse pre-implantation embryo development

Changes in metabolite levels during early embryogenesis are presented as a heatmap in Fig. 1e. The dynamics of some metabolites were consistent with previous studies, including α-ketoglutarate (α-KG), succinate, and malic acid, which are higher in the later stages, and nicotinamide, NAD^+^, AMP and glutathione, which are higher in the earlier stages.^17^ Furthermore, metabolomic profiles were mapped to their respective biochemical pathways using the Kyoto Encyclopedia of Genes and Genomes (KEGG), revealing distinct metabolite patterns (Fig. 1f). Nucleic acid is an important class of macromolecules found in all cells; it is produced in different molecular forms that participate in protein synthesis. Several key intermediates of nucleic acid metabolism exhibited high levels from the 4-cell stage, such as purine, adenine, and uracil (Fig. 1e). KEGG analysis also revealed pentose phosphate (PPP) and pyrimidine/purine metabolism intermediates were enriched from the 4-cell embryo to blastocyst. Amino acids are intermediates in nucleotide and lipid biosynthesis, carbon and nitrogen homeostasis, and redox buffering. Our temporal metabolite profiles showed variation in the levels of certain amino acids during early embryogenesis. We detected the specific abundance of some amino acids from the 8-cell embryo to blastocyst stages, such as glutamine in the 8-cell embryo, methionine in the morula, and lysine and serine in the blastocyst (Fig. 1e). KEGG analysis revealed that alanine, aspartate (Asp), and glutamate metabolism were upregulated during these processes (Fig. 1f). These diverse metabolic trends in early embryos reflect the complex, fine-tuned regulation of amino acid metabolism. It has been evidenced that carbohydrate metabolism activity is lower in early cleavage embryos than blastocysts and somatic cells.^16,18^ TCA cycle metabolites such as aconitic acid, α-KG, succinate, malic acid, and oxaloacetate were elevated during development from the 8-cell embryo to blastocyst (Fig. 1e); KEGG analysis also revealed the enrichment of TCA cycle during the later stages (Fig. 1f). To verify our metabolomic data, we further performed targeted metabolomic analysis of carbohydrate metabolism using 24,000 embryos from the zygote to blastocyst stages, each with four biological replicates (Supplementary Table S2). The levels of 11 metabolites were quantified (Supplementary Fig. S1a); their dynamics were similar to those of untargeted metabolites, such as NAD^+^, NADP^+^, citric acid and oxaloacetate, indicating high confidence of metabolomic data with our approach (Supplementary Fig. S1b). Taken together, these data on metabolome dynamics during mouse pre-implantation embryo development provide a reference such that stage-specific metabolite networks may be used to define developmental progress.

### Activities of metabolic pathways during mouse pre-implantation embryo development

To better understand the involvement of metabolic pathways in mouse early development, we mapped all enzymes, which are mediators of cellular metabolism, from transcriptomic and proteomic data on mouse early embryos to the main KEGG metabolic pathways associated with the metabolites (Fig. 2a and Supplementary Fig. S2a).^19,20^ Substantial evidence indicates that TCA cycle activity occurs at the later stage of early embryos.^17^ Consistent with this notion, the expression of TCA cycle enzymes was upregulated from 8-cell embryo and peaked in blastocyst. Also, the enrichment of the TCA cycle key metabolites (aconitic acid, isocitric acid, α-KG, succinate, malic acid, oxaloacetate) were elevated from 8-cell embryo to blastocyst, showing the enhanced activity of TCA cycle from 8-cell embryo and peaks at the blastocyst. It should be noted that citric acid was fell in the blastocyst, which may be due to the high transition from citric acid to acetyl-CoA, supporting by the high expression level of *Acly* in the blastocyst. Three intermediates associated with glycolysis (glucose, F-1,6-BP and pyruvic acid) we identified began to be accumulated in the blastocyst. Meanwhile, the expression of glycolytic enzymes underwent a dramatic increase at the blastocyst stage (such as *Hk, Gpi, Aldo, Tpi, Gapdh, Pgk, Pgam, Eno* and *Pkm*), suggesting the activation of glycolysis in the blastocyst. In addition, glutamine peaked in the 8-cell embryo, and the integrated analysis of proteomic and transcriptomic data showed remarkably elevated levels of *Glud1* was observed in the blastocyst, suggesting the activation of glutaminolysis metabolism, which contribute to produce α-KG from glutamate, happens at the blastocyst stage. We also found that methionine and S-adenosylhomocysteine associated with methionine cycle were elevated at the morula stage,^21^ and the proteomic and transcriptomic data showed reproducible and statistically significant upregulation of *Mat2a* level after 8-cell embryo, suggesting the activation of methionine cycle from 8-cell embryo to blastocyst. To define the transition of stage-specific transcriptome-metabolome networks during the developmental process, we performed weighted gene co-expression network analysis (WGCNA) using the top 5,000 genes and high metabolites at each stage; a total of nine co-expression modules were identified (Fig. 2b; Pearson’s correlation coefficient ≥ 0.6, P < 0.05). Among these modules, we focused on that specific to the 2-cell embryo, where ZGA occurs. Genes in this module were involved in ZGA genes, such as *Zscan5b, Zscan4d* and *Arl16*, and metabolites, including lactate, nicotinamide ribotide and NAD^+^, were observed (Supplementary Fig. S1c). To further screen functional metabolites associated with ZGA, we conducted correlation analysis of each metabolite with other metabolites and the top 5,000 genes at the 2-cell stage. The results revealed close connections between NAD^+^ and both 2-cell genes and metabolites (Fig. 2c), suggesting that it has a functional role in ZGA.

**Fig. 2.**
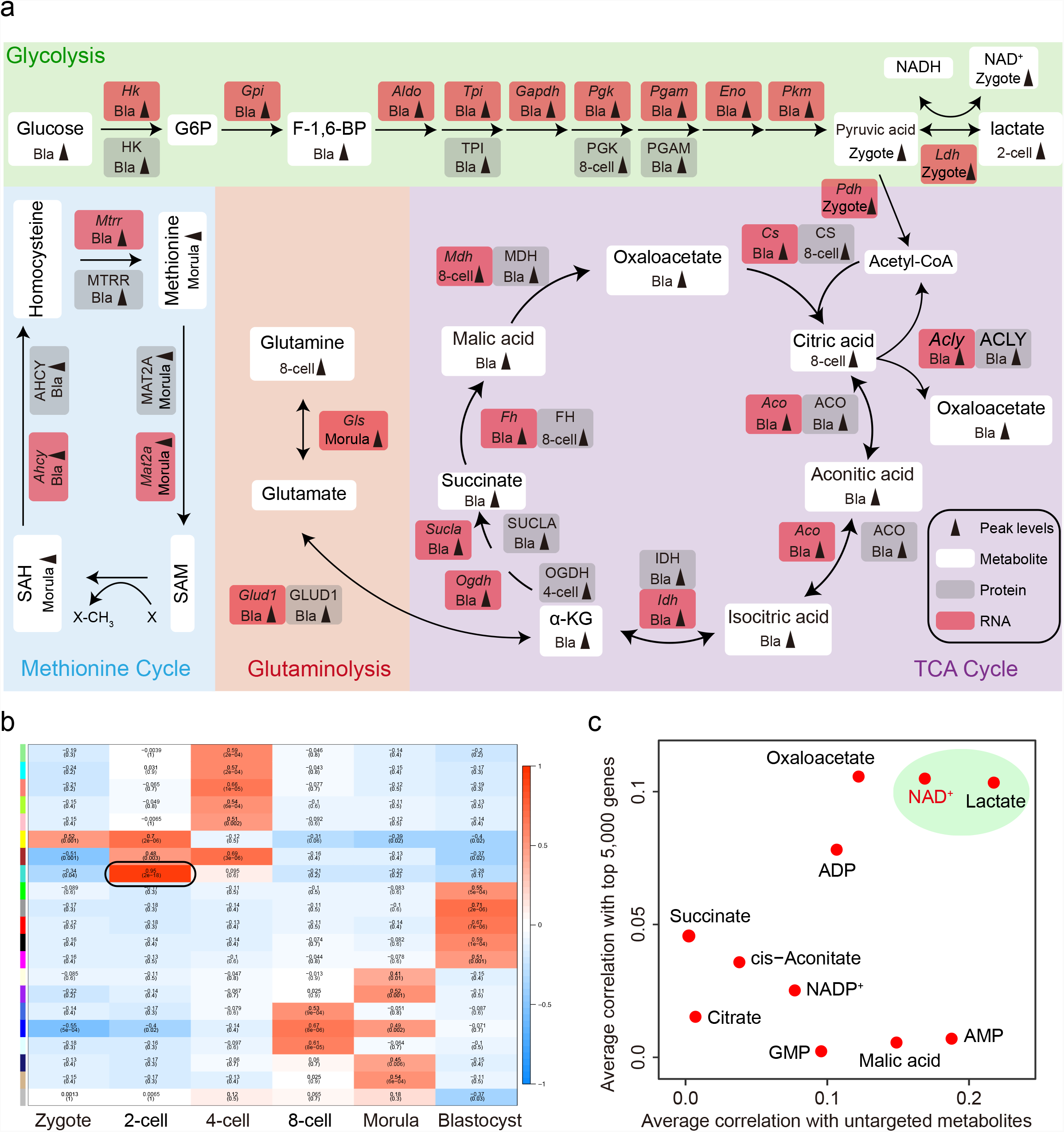
Activities of metabolic pathways during mouse pre-implantation embryo development. **a** Schematic diagram of carbohydrate metabolism, integrated with transcriptomic and proteomic data. Grey arrows indicate peak level at indicated stage. White boxes represent metabolites, and gray and red boxes represent protein and mRNA of indicated metabolic enzymes, respectively. **b** Stage-specific co-expression genes and metabolite modules and their correlation to development stage. Each row corresponds to a co-expression module. Numbers in each square represent correlation coefficients between the module and development stage, with *p*-values in brackets. Red and blue squares indicate positive and negative correlations, respectively; white indicates no correlation. **c** Correlation of metabolites and genes at the 2-cell embryo stage. The X-axis shows the correlations of a specific metabolite with the top 5,000 genes. The Y-axis shows the correlations of a specific metabolite with other metabolites.

### NAD^+^ synthesis is indispensable for mouse embryo development

NAD^+^ is a key cofactor for hundreds of processes related to energy metabolism.^22^ Therefore, we experimentally manipulated its function during early embryogenesis. In bioenergetic pathways, NAD is reversibly converted between NAD^+^ and its reduced (NADH) state. First, we determined the NAD^+^ and NADH levels during mouse early embryo development. Consistent with the large consumption of NAD^+^ reported previously,^23^ NAD^+^ level decreased significantly from the zygote to 2-cell embryo stages, the remarkable decline of NADH was observed at the 8-cell stage, and increased at the blastocyst stage (Fig. 3a). De novo biosynthesis of NAD^+^ is from tryptophan (Trp) by the kynurenine pathway. To explore the role of NAD^+^ in mouse embryo development, we deprived of Trp from the culture medium, but the development of embryos cultured in the medium was not affected (Supplementary Fig. S3a). The nicotinamide (NAM) salvage pathway is the predominant route for mammalian NAD^+^ biosynthesis (Fig. 3b). We treated zygotes with FK866, an inhibitor of the NAD^+^ biosynthetic rate-limiting enzyme NAM phosphoribosyltransferase (*Nampt*), whose RNA level was high from the middle 2-cell to 4-cell embryos and in blastocyst, and in consistent with that the protein level was elevated from 2-cell embryo and reached to the peak in blastocyst, indicating the activity of NAM salvage pathway at the early stage of mouse embryonic development (Supplementary Fig. S3b). As shown in Fig. 3c, FK866-treated embryos exhibited complete developmental arrest from the 4- to 8-cell stages, and nicotinamide mononucleotide (NMN) treatment fully rescued developmental arrest, suggesting the importance of NAD^+^ synthesis in early embryo development (Fig. 3c-e and Supplementary Fig. S3c, d). In the cytosol, the interconversion of pyruvic acid and lactate is generally thought to be in equilibrium with free NAD^+^ and NADH through the catalytic action of lactate dehydrogenase (LDH) (Fig. 3f).^24^ To assess the contribution of this transition to NAD^+^ biosynthesis, we treated zygotes with GNE-140, which is a dual LDHA/B inhibitor, resulting in a development block from the 8- to 16-cell stages, which was rescued by NMN addition (Fig. 3g-i and Supplementary Fig. S3e). We complemented this approach using the LDHA-specific inhibitor oxamate. Unexpectedly, embryo development was not affected by oxamate (Fig. 3g, h and Supplementary Fig. S3f), suggesting that LDHB, but not LDHA, may contribute to NAD^+^ production, consistent with the expression patterns showing that only *Ldhb* was highly expressed at the earlier stage (Supplementary Fig. S3g, h). Asp plays an important role in the malate-aspartate (Mal-Asp) shuttle, which controls the NAD^+^/NADH ratio in cytoplasm (Fig. 3j).^16^ However, the development of embryos cultured in medium lacking Asp was not affected (Fig. 3k-m), and Asp addition did not rescue the deficiencies caused by FK866 and GNE-140 treatment (Fig. 3n), indicating that the Mal-Asp shuttle little affects the NAD^+^ levels of 2-cell embryos. Consistent with these results, the NAD^+^ level decreased significantly in FK866- and GNE-140-treated embryos, and was rescued by NMN addition, but was not affected by Asp deficiency (Fig. 3o). Considering the dramatic changes of NAD^+^ level at the 2-cell stage, and that developmental arrest in embryos deficient with NAD^+^ synthesis occurred first in the 4-cell stage, we propose that NAD^+^ mainly functions between the 2- and 4-cell stages. Therefore, we switched the medium between control and FK866 addition at the 2- and 4-cell stages. As expected, the deprivation of NAD^+^ from 2- to 4-cell embryos resulted in failure to develop to the blastocyst stage (Fig. 3p-r). Outside of this temporal window, embryo development was not affected. Together, these results demonstrate that NAD^+^ synthesis is important for the development of mouse embryos from the 2- to 4-cell stages, suggesting large amount of NAD^+^ is consumed at the stages.

**Fig. 3.**
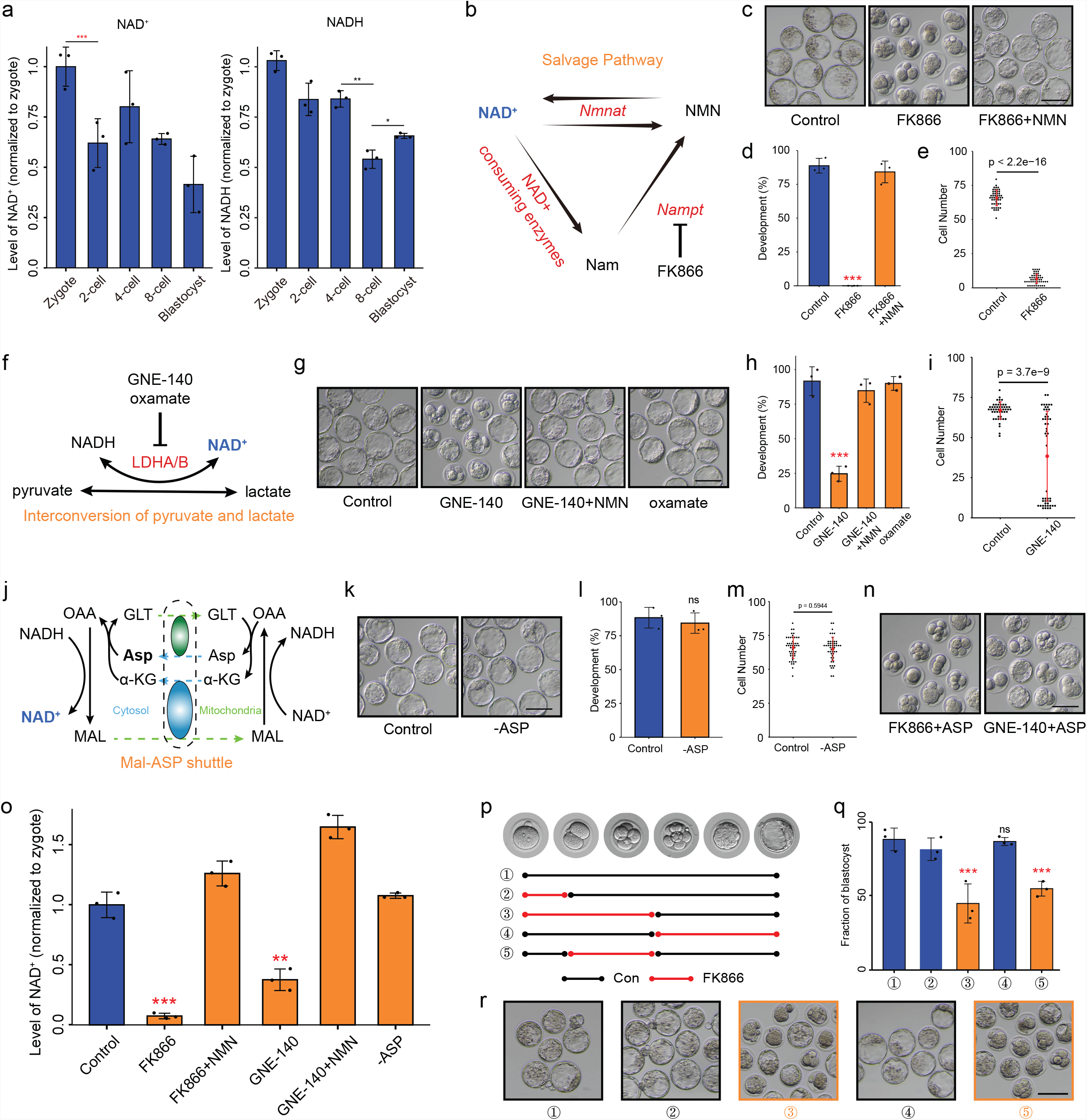
NAD^+^ depletion impairs early mouse embryo development. **a** NAD^+^ and NADH levels during early mouse embryo development. **b** Overview of the NAM salvage pathway. **c** Representative images and (**d**) developmental rates of control, FK866-treated, and FK866+NMN-supplemented embryo groups. Data are from three independent experiments. ***P < 0.001 (Student’s *t*-test). Scale bars, 50 μm. **e** Quantification of cells in control and FK866-treated embryo groups. Dots represent cell numbers of single embryos. **f** Overview of the interconversion pathway of pyruvic acid and lactate. **g** Representative images and (**h**) developmental rates of control, GNE-140-treated, GNE-140+NMN-supplemented, and oxamate-treated embryos. Data are from three independent experiments. ***P < 0.001 (Student’s *t*-test). Scale bars, 50 μm. **i** Quantification of cells in control and GNE-140-treated embryos. Dots represent cell numbers of single embryos. **j** Overview of the Mal-Asp shuttle. GLT: glutamate; MAL: malate; OAA: oxaloacetate. **k** Representative images and (**l**) developmental rates of control and Asp-deletion embryos. Data are from three independent experiments. ***P < 0.001 (Student’s *t*-test). Scale bars, 50 μm. **m** Quantification of cells in control and Asp-deletion embryos. Dots represent cell numbers of single embryos. **n** Development of FK866- and GNE-140-treated embryos supplemented with Asp. **o** NAD^+^ level in control, FK866-treated, FK866+NMN, GNE-140-treated, GNE-140+NMN, and Asp-deletion late 2-cell embryos. **p-r** Timeline of FK866 treatment. Embryos were cultured in control or FK866-addition medium. The stages of FK866 treatment are indicated in (**p**). Developmental rates (**q**) and representative images (**r**) of each group are shown. Data are from three independent experiments. **P < 0.01, ***P < 0.001 (Student’s *t*-test). Scale bars, 50 μm.

### Depletion of NAD^+^ synthesis disturbs minor ZGA in early mouse embryos

To further investigate the mechanism by which NAD^+^ synthesis influences early mouse embryo development, we performed RNA sequencing (RNA-seq) of late 2- and 4-cell embryos with and without FK866 treatment. Unsupervised hierarchical clustering (UHC) was applied to separate FK866-treated and control embryos (Supplementary Fig. S4a). A total of 1,494 and 1,248 significantly differentially expressed genes (DEGs) were detected between the control and FK866-treated groups among late 2- and 4-cell embryos, respectively (P < 0.05, Fig. 4a, b and Supplementary Table S3). Notably, most genes upregulated upon FK866 treatment in the late 2- and 4-cell embryos were ZGA genes, and their associated pathways were related to ZGA (Fig. 4c and Supplementary Fig. S4b, c). A previous study classified ZGA genes into minor and major ZGA genes based on their expression, mainly at the early or late 2-cell stages.^25^ According to this classification, we compared the effect of NAD^+^ depletion on minor and major ZGA. In late 2-cell embryos, 32.7% of genes upregulated by FK866 treatment were minor ZGA genes; the ratio reached 21.6% in 4-cell embryos, indicating severe impairment of the minor ZGA (Fig. 4d). We also found that *Duxf3* (*Dux*), a master transcription factor that transiently regulates the ZGA process and is specifically expressed in early mouse 2-cell embryo,^26^ was significantly upregulated in the FK866-treated late 2-cell embryo (P = 0.0095, log_2_fold change = 4.87; Fig. 4a). Overexpression of *Dux* was confirmed by quantitative polymerase chain reaction (qPCR) and immunostaining (Fig. 4e-g). These results demonstrate that NAD^+^ deprivation leads to prolonged expression of many minor ZGA genes until the late 2- and 4-cell stages. Next, we investigated whether NMN could rescue excessive ZGA. All minor ZGA genes (n=1,065), which transiently expressed at the late zygote and early 2-cell stages, were clustered into four groups. Cluster 1 and cluster 2 genes were upregulated by FK866 treatment, of which NMN recovered the overexpression of 62% and 72.2% genes (cluster 1) in late 2- and 4-cell embryos, respectively, including *Dux* (Fig. 4h and Supplementary Fig. S4d, e and Table S4), indicating a significant role of NAD^+^ synthesis in moderate minor ZGA timely. Aberrant minor ZGA, such as the prolonged activation of *Dux*, severely impairs major ZGA and early embryo development.^6^ Therefore, these results indicate that the failure of early development upon NAD^+^ deprivation is at least partly caused by excessive minor ZGA.

**Fig. 4.**
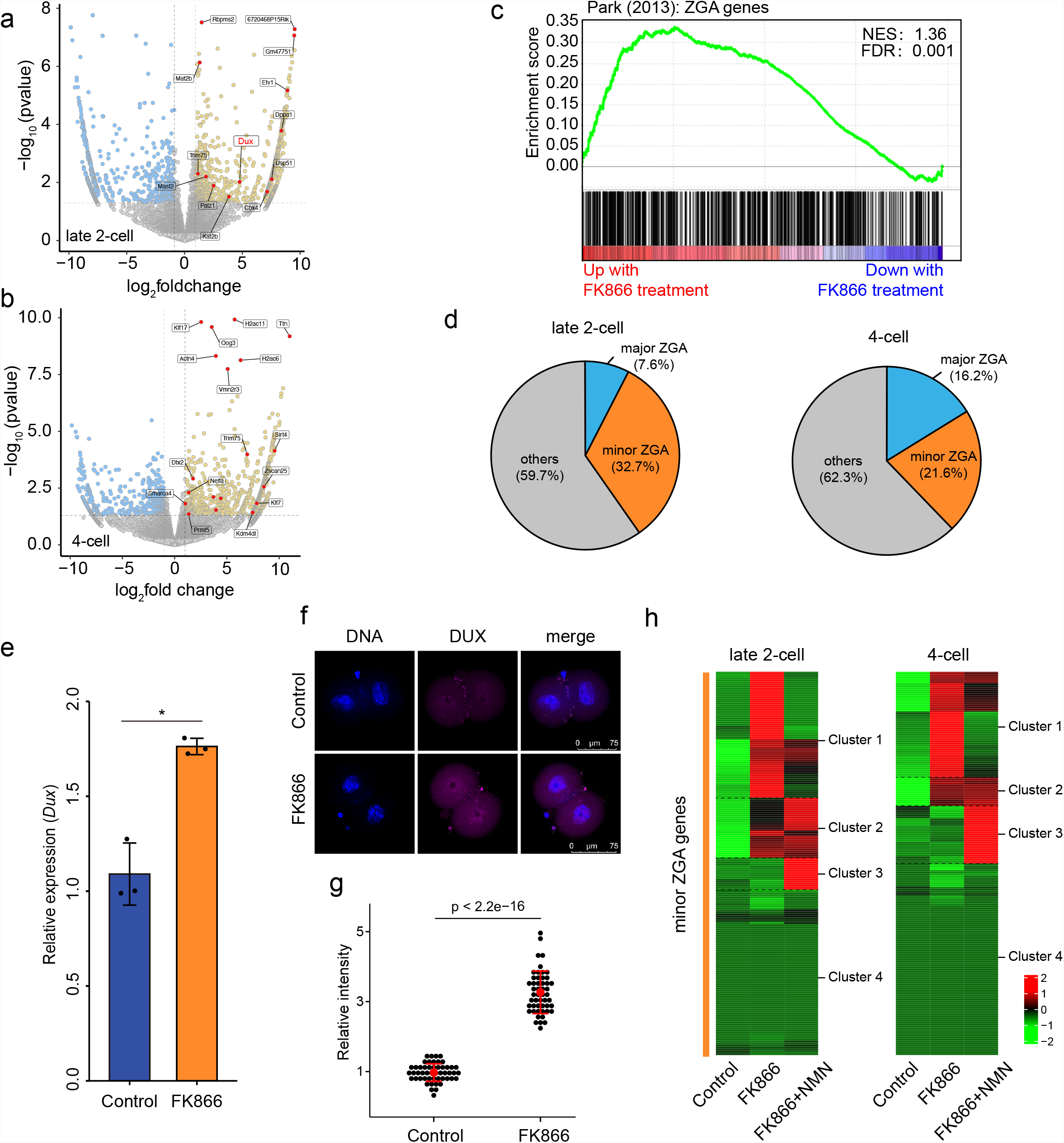
NAD^+^ depletion disturbs minor ZGA in early mouse embryos. RNA-seq analysis of control and FK866-treated (**a**) late 2-cell and (**b**) 4-cell mouse embryos. Volcano plots show gene expression changes. Yellow and blue dots indicate upregulated (fold change > 1) and downregulated (fold change < −1) genes with P < 0.05. **c** Gene set enrichment analysis (GSEA) of ZGA genes showed preferential upregulation in FK866-treated late 2-cell embryos. **d** Pie chart showing the fraction of minor and major ZGA genes upregulated in late 2-cell and 4-cell embryos. **e** Results of quantitative polymerase chain reaction (qPCR) showing *Dux* upregulation in FK866-treated late 2-cell embryos. Data are means ± standard error of the mean (SEM) from three independent experiments. **f** Representative confocal images of control and FK866-treated embryos stained with *Dux* antibody. Scale bars, 75 µm. **g** Quantification of DUX fluorescence intensity in control and FK866-treated embryos. Each dot represents a single nucleus. **h** Heatmaps show that minor ZGA genes were upregulated between control and FK866-treated embryos, which was rescued by NMN supplementation. NMN recovered the overexpression of 62% and 72.2% minor ZGA genes at late 2-cell and 4-cell stages, respectively. Mean values of two biological replicates were scaled and are represented as Z scores.

### NAD^+^ synthesis is important for zygotic H3K27ac erasure

Previous studies have shown that NAD^+^ functions as an obligated cofactor for histone deacetylation,^22^ which is related to chromatin accessibility.^27^ We therefore questioned whether the change in the histone acetylation mediated by NAD^+^ account for the regulation of minor ZGA. Among the most abundant histone acetylation modification, H3K9ac and H3K27ac play a substantial role in defining chromatin states and modulating gene expression.^28^ In agreement with previous study,^29^ H3K9ac showed no remarkable changes during mouse preimplantation development (Supplementary Fig. S5a, b). However, H3K27ac was dramatically changed (Fig. 5a, b). In contrast to the weak signals in MII oocytes and sperm, we observed a strong increase in H3K27ac levels at the zygote stage. H3K27ac signals were rapidly lost from the middle to late 2-cell stages, and increased progressively after the 4-cell stage (Fig. 5a, b). To further determine when H3K27ac is established after fertilization, we evaluated the dynamics of H3K27ac across all pronuclear stages of the zygote. The results showed that H3K27ac was established as early as pronuclear stage 1 (PN1) (Supplementary Fig. S5c, d), suggesting that H3K27ac may participate in the initiation of the early development process. Then, we used cleavage under targets and tagmentation (CUT&Tag) to generate genome-wide H3K27ac histone modification maps for mouse sperm, oocyte, zygote (PN4), early 2-cell, late 2-cell, and 4-cell embryos (Supplementary Fig. S5e). Genomic coverage analysis revealed 75,496 peaks of established H3K27ac modifications in the zygote; these peaks decreased rapidly in the late 2-cell embryo (Supplementary Fig. S5e, f). We annotated the distribution of the zygote-established H3K27ac domains (zyH3K27ac) through model-based analysis of chromatin immunoprecipitation assay sequencing (ChIP-seq) (MACS), and found that they were mainly enriched in the promoter, intron, and intergenic regions (Supplementary Fig. S5g). Then, we summarized the zyH3K27ac results according to three chromatin states based on their normalized levels, and found that the dynamics of zyH3K27ac were clearly highly established in the zygote, maintained in the early 2-cell embryo, and lost in the late 2- and 4-cell embryos (Fig. 5c). The enrichment of zyH3K27ac temporally precedes minor ZGA, implying a role for zyH3K27ac in minor ZGA regulation. Next, we tested the role of NAD^+^ synthesis in the dynamics of H3K27ac modifications in early mouse embryos using immunofluorescence staining. FK866-treated embryos retained high levels of H3K27ac at the late 2- and 4-cell stages, which was rescued by supplementation with NMN (Fig. 5d, e and Supplementary Fig. S5h, i); this indicated a significant role of NAD^+^ synthesis in the deacetylation of H3K27ac in mouse cleavage embryos. Analysis of CUT&Tag data for control, FK866, and NMN treated-embryos showed that 52.5% of zyH3K27ac deacetylation peaks in the control remained hyperacetylated in FK866-treated embryos, and were eliminated by NMN treatment in the late 2-cell embryo (Fig. 5f, g); a similar phenomenon was observed in the 4-cell embryo (Supplementary Fig. S5j). These results suggest that NAD^+^ synthesis is important for zyH3K27ac erasure at late 2-cell stage.

**Fig. 5.**
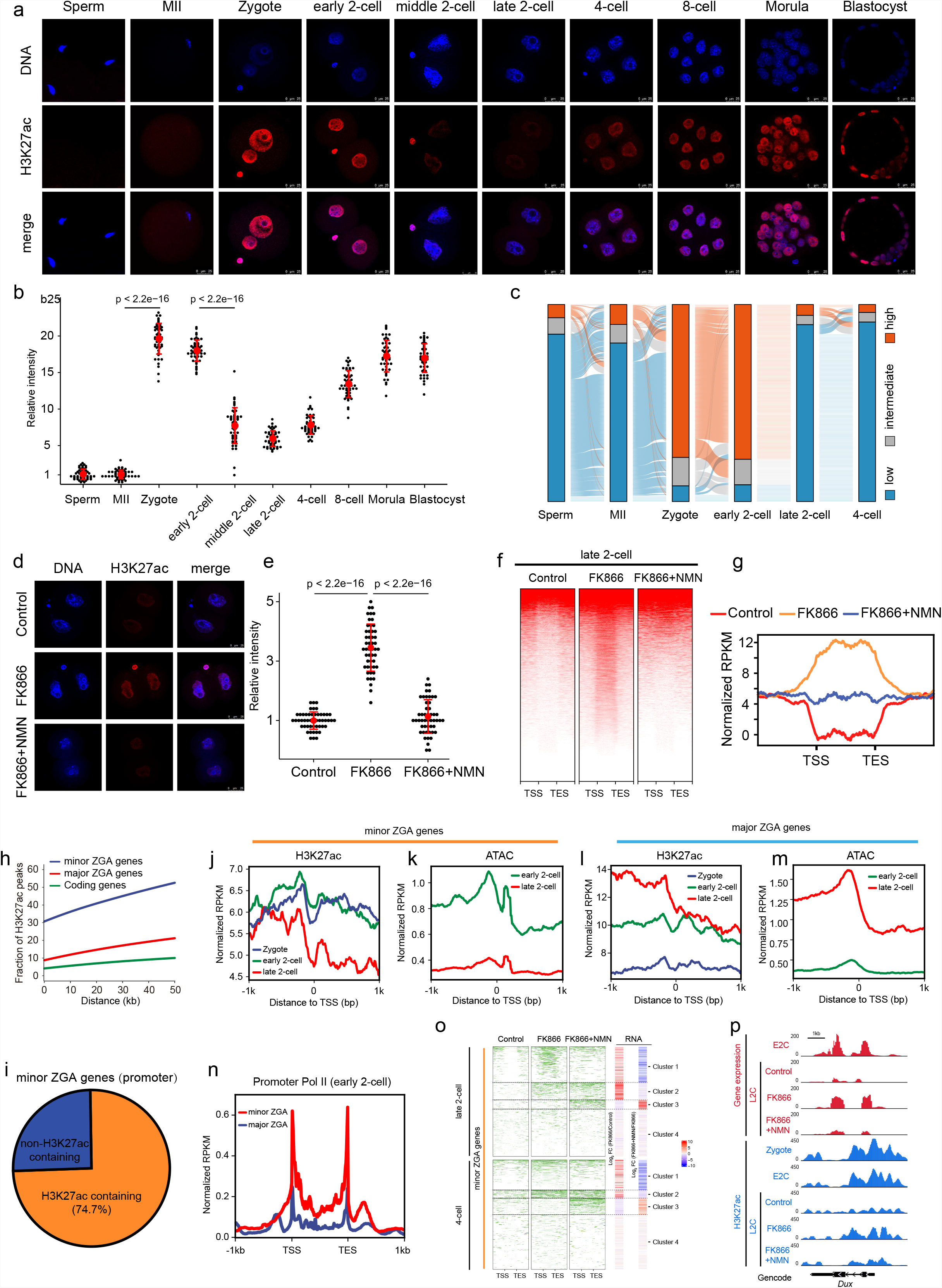
NAD^+^-mediated erasure of zygotic H3K27ac avoids excessive minor ZGA. **a** Immunostaining of H3K27ac during mouse pre-implantation embryo development. One representative image from three independent experiments is shown. Scale bar, 25 μm. **b** Quantification of H3K27ac fluorescence intensity in mouse embryos. Each dot represents a single nucleus. **c** Alluvial plots showing the global dynamics of zyH3K27ac during early embryo development. **d** Representative confocal images of control, FK866-treated, and FK866+NMN-treated late 2-cell embryos stained with H3K27ac antibody. Scale bars, 25 µm. **e** Quantification of H3K27ac fluorescence intensity in late 2-cell embryos. Each dot represents a single nucleus. **f** Heatmap showing H3K27ac signals ranked by their relative change after FK866 treatment. NMN rescued the changes induced by FK866 treatment in late 2-cell embryos. **g** Metaplot of H3K27ac peaks (Z-score normalized; n = 15,920) in control, FK866-treated, and FK866+NMN-treated late 2-cell embryos. **h** Comparison of the distance between H3K27ac peaks and ZGA or coding genes in the genome. The Y-axis shows the proportion of H3K27ac peaks associated with genes. About 30% to 50% of H3K27ac peaks are distributed from the TSSs to a distance of 50 kb to the TSSs of minor ZGA genes, and the proportions are higher than those of major ZGA genes and other coding genes. **i** Pie charts showing the percentages of minor ZGA gene whose promoters enriched the zyH3K27ac at the zygote stage. Metaplot showing the H3K27ac enrichment (Z-score normalized) at the promoters of the minor ZGA genes (**j**) and major ZGA genes (**l**) at individual stages. Metaplot showing the ATAC-seq enrichment (Z-score normalized) at the promoters of the minor ZGA genes (**k**) and major ZGA genes (**m**) at individual stages. **n** Metaplot showing the Pol II enrichment (Z-score normalized) at the promoters of minor and major ZGA genes at the early 2-cell stages. **o** Heatmap (left) showing H3K27ac signals ranked by their relative changes following FK866 or FK866+NMN treatment. Heatmap (right) shows that minor ZGA genes were upregulated between control and FK866-treated embryos, and rescued by NMN supplementation. Mean values of two biological replicates were scaled and are represented as Z scores. **p** Genome browser view of RNA-seq and H3K27ac signals at the *Dux* locus in control, FK866-treated, and FK866+NMN-treated late 2-cell embryos.

### NAD^+^-mediated erasure of zygotic H3K27ac is required for the exit of minor ZGA

The dynamic of zyH3K27ac shows its possible role in minor ZGA regulation. We found that, within a distance of 50 kb, the distribution of the zyH3K27ac peaks that neighbored minor ZGA genes was much higher than that of major ZGA genes and other coding genes (Fig. 5h; Student’s *t*-test, P < 8.15 × 10^−4^), raising the possibility that the activation of minor ZGA genes may be driven by zyH3K27ac. Actually, 74.7% of minor ZGA gene promoters enriched zyH3K27ac signals (Fig. 5i), and these signals were high in the early 2-cell embryo, and low in the late 2-cell embryo (Fig. 5j and Supplementary Fig. S5k). H3K27Ac reflects a loosened chromatin state.^30^ In line with the dynamic of zyH3K27ac at the minor ZGA gene promoters, we found that the promoters were open at the early 2-cell stage, and close at the late 2-cell stage (Fig. 5k and Supplementary Fig. S5l). Whereas, the zyH3K27ac and ATAC signals in major ZGA gene promoters were high at the late 2-cell stage (Fig. 5l, m and Supplementary Fig. S5m, n). The relationship between the enrichment of zyH3K27ac and the expression of minor ZGA genes was further confirmed by Pol II small-scale Tn5-assisted chromatin cleavage with sequencing (Stacc–seq) data.^31^ At the early 2-cell stage when the zyH3K27ac and ATAC signals were enriched in the promoters of minor ZGA genes but not major ZGA genes, Pol II showed higher enrichment at minor ZGA genes than major ZGA genes (Fig. 5n). These observations are in agreement with the notion that zyH3K27ac may drive minor ZGA by establishing chromatin accessibility. We further address the role of NAD^+^-mediated erasure of zyH3K27ac in regulating minor ZGA. We found that changes in the expression of minor ZGA genes and their associated zyH3K27ac peaks were highly correlated among the control, FK866-treated, and NMN-supplemented groups (Fig. 5o). Notably, the zyH3K27ac peaks associated with genes in cluster 1, including *Dux* (Fig. 5p), showed high by FK866 treatment and recovered by NMN supplement, which was consistent with the upregulated and downregulated expression (Fig. 5o).

To demonstrate the effect of zyH3K27ac erasure on minor ZGA and early embryo development, we treated mouse embryos from the early 2- to 4-cell stages using trichostatin A (TSA) to inhibit the erasure of zyH3K27ac (Supplementary Fig. S6a). We found that the development of embryos treated with 10 nM TSA was arrested mainly at the 4-cell stage (Supplementary Fig. S6b-d). As expected, TSA-treated embryos showed high H3K27ac levels at the late 2-cell stage (Supplementary Fig. S6e, f). Consistently, CUT&Tag data revealed that nearly half of the zyH3K27ac peaks (32,251 of 75,496) remained hyperacetylated after TSA treatment (Supplementary Fig. S6g, h). we performed RNA-seq on late 2-cell embryos with and without TSA treatment. UHC was performed to separate TSA-treated embryos from control embryos (Supplementary Fig. S6i). A total of 2,257 and 1,492 genes were significantly up- or downregulated between the control and TSA-treated late 2-cell embryos, respectively (P < 0.05; Supplementary Fig. S6j and Table S5). The expression of many minor ZGA genes was found to be dramatically elevated compared to the control group, and their associated zyH3K27ac peaks remained high after TSA treatment (Supplementary Fig. S6k). These results suggest that NAD^+^-mediated erasure of zyH3K27ac contributes to the precise and timely regulation of ZGA at the late 2- and 4-cell stages.

### NAD^+^ could cooperate with *Sirt1* to remove zyH3K27ac

Sirtuins, including SIRT1-7, are a family of NAD^+^-dependent protein deacetylases.^22^ To explore the role of the candidate histone deacetylase, which is responsible for zyH3K27ac erasure in early mouse embryos, we examined the expression patterns of SIRTs. RNA-seq data revealed that only *Sirt1* was highly expressed in middle and late 2-cell embryos (Supplementary Fig. S7a, b). This expression pattern was confirmed by qPCR, and immunostaining showed that the SIRT1 protein was highly expressed in the nuclei of embryos from the middle 2- to 4-cell stages (Fig. 6a-c), coincident with the timing of zyH3K27ac erasure. This expression pattern prompted us to investigate whether *Sirt1* plays an important role in deacetylating zyH3K27ac. We knocked down *Sirt1* at the RNA and protein levels by injecting siRNA, *Trim21* mRNA, and SIRT1 antibody into zygotes (Fig. 6d). Successful knockdown was confirmed by qPCR and immunostaining (Supplementary Fig. S7c-e). Compared to the control group, the potential to develop to the blastocyst stage in *Sirt1*-depleted embryos was reduced (90% vs. 67.3%, P < 0.005) and blocked mainly at the morula stage, and the NMN addition did not rescue the failure of blastocyst formation, suggesting that NAD^+^ depends on *Sirt1* to remove H3K27ac (Fig. 6e, f). Definitely, embryos showing *Sirt1* deficiency retained high levels of H3K27ac modifications at the late 2-cell stage (Fig. 6g, h), but the H3K9ac levels were not significantly affected (Supplementary Fig. S7f), suggesting *Sirt1* facilitates the deacetylation of H3K27ac, but not H3K9ac, during the process of ZGA. Furthermore, about 30% of zyH3K27ac peaks were sustained after *Sirt1* knockdown according to CUT&Tag data analysis (Fig. 6i and Supplementary Fig. S7g). RNA-seq analysis revealed that 690 and 659 DEGs were up- and downregulated, respectively, in *Sirt1*-knockdown late 2-cell embryos compared to control embryos (Supplementary Fig. S7h, i and Table S6). We also observed a tendency of minor ZGA genes to show higher expression in *Sirt1*-depleted embryos, which was highly correlated with the failure of zyH3K27ac removal (Supplementary Fig. S7j). These results support the notion that SIRT1 is highly expressed at the middle 2-cell stage, and contributes to NAD^+^-mediated zyH3K27ac deacetylation.

**Fig. 6.**
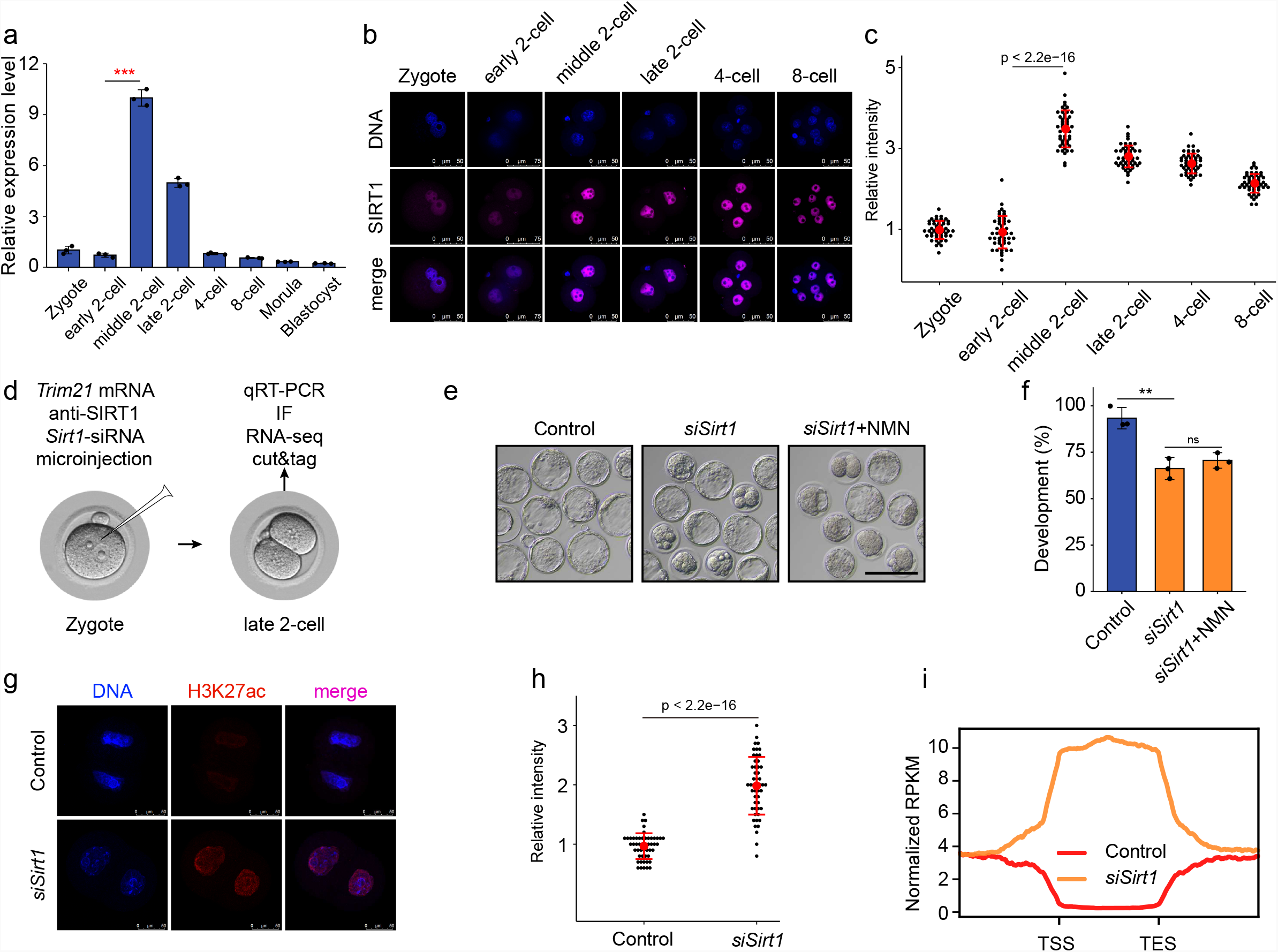
NAD^+^ could cooperate with *Sirt1* to remove zyH3K27ac. **a** The relative expression of *Sirt1* was measured by quantitative reverse-transcribed PCR (qRT-PCR) analysis. Error bars indicate standard deviation (SD). **b** Immunostaining of SIRT1 during mouse pre-implantation embryo development. One representative image from three independent experiments is shown. Scale bar, 50 μm. **c** Quantification of SIRT1 fluorescence intensity in mouse embryos. Each dot represents a single nucleus. **d** Schematic presentation of the *Sirt1* knockdown experimental protocol. **e** Representative images and (**f**) developmental rates of control and *Sirt1* knockdown embryos. NMN (10 µM) addition did not rescue blastocyst formation of *Sirt1*-knockdown embryos. Data are from three independent experiments. **P < 0.01 (Student’s *t*-test). Scale bars, 50 μm. **g** Representative confocal images of control and *Sirt1*-knockdown embryos stained with H3K27ac antibody. Scale bars, 75 µm. **h** Quantification of H3K27ac fluorescence intensity in control and *Sirt1* knockdown embryos. Each dot represents a single nucleus. **I** Metaplot of H3K27ac signals (Z-score normalized) in control and *Sirt1* knockdown late 2-cell embryos.

### NAD^+^ supplement promotes the removal of zygotic H3K27ac and benefits preimplantation development of human embryos

The finding that NAD^+^ is crucial for timely modulation of minor ZGA and further embryonic development of mouse embryo led us to investigate whether a similar mechanism might operate in human embryos. In human, minor ZGA occurs at the 2- to 4-cell stages, and major ZGA activates at the 4- to 8-cell stages.^32^ We performed immunostaining of H3K27ac in human embryos. The high H3K27ac signals were observed in the zygote and 2-cell embryo, and remarkably reduced H3K27ac in the 8-cell embryo (Fig. 7a). The finding shows that, similar to that seen in mice, H3K27ac deacetylation occurs when major ZGA activates in human embryo. Furthermore, RNA-seq data revealed that *SIRT1* was dramatically elevated in the 8-cell embryo (Fig. 7b), and *SIRT1* proteins are conserved between human and mouse (Supplementary Fig. S7k). The results presume that *SIRT1* cooperated with NAD^+^ to remove zyH3K27ac ensures the timely silencing of minor ZGA program in human embryo. To address the notion, we treated human zygotes by NMN addition. As expected, NMN-treated group showed weaker signals compared to control group at 8-cell stage (Fig. 7a). Importantly, more the NMN-treated embryos could develop to blastocyst than the control group (Fig. 7c; 36.4% VS. 11.1%), and the developmental process was normal (Fig. 7d and Supplementary Movie S1). Taken together, the mechanism of NAD^+^-SIRT1 -mediated deacetylation may share between mice and human, which is crucial for mammalian embryo development.

**Fig. 7.**
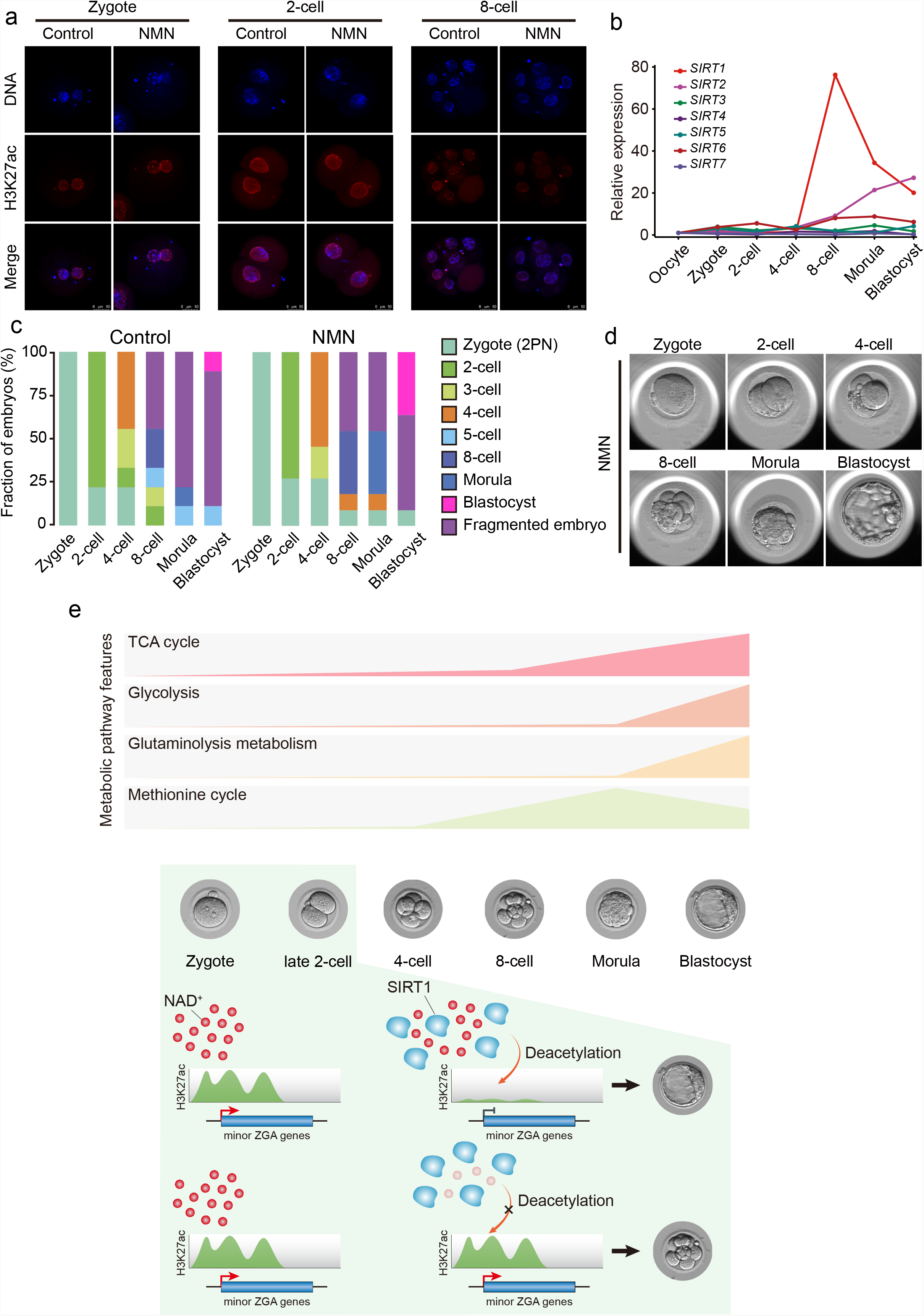
NAD^+^ supplement promotes the removal of zygotic H3K27ac and benefits preimplantation development of human embryos. **a** Representative confocal images of control and NMN-treated human embryos stained with H3K27ac antibody. Scale bars, 50 µm. **b** Sirtuin family expression patterns according to RNA-seq data in human pre-implantation embryos. **c** Stacked bar plots show fraction of human embryos at the different developmental stages after NMN addition (right side, n = 11), or control (left side, n = 9). **d** Representative images of the development of NMN-treated human embryos. **e** Metabolic characteristics during mouse pre-implantation development (upper) and a model of NAD^+^ involvement in the precise and timely regulation of minor ZGA (below).

## Discussion

Decades of research have highlighted the important role of cell metabolism in mammalian pre-implantation development. However, due to limited amounts of experimental material, a comprehensive investigation of metabolite dynamics during the process is lacking. In this study, we established global metabolomic profiles of mouse pre-implantation embryos from the zygote to blastocyst stages. Using functional approaches, we further demonstrated the important role of NAD^+^ synthesis in the precise and timely regulation of minor ZGA, which is indispensable for mouse pre-implantation development (Fig. 7e).

The metabolism of mammalian early embryos has been investigated mainly using radiolabeled substrates in combination with inhibitors, and exogenous substrates at specific stages.^33^ For example, embryos utilize pyruvic acid early, and later transition to glucose.^34^ Further study is required to fully understand the dynamics of endogenous metabolites during the entire mammalian pre-implantation development process. In this study, to precisely establish the metabolic landscape, we collected thousands of mouse embryos at all pre-implantation stages from zygote to blastocyst, and examined the dynamics of hundreds of metabolites related to amino acid, carbohydrate, lipid, and nucleotide metabolism. We integrated these data with transcriptomic and proteomic data, and discovered the metabolic characteristics of the development process, including the activation of methionine cycle from 8-cell embryo to blastocyst, high glutaminolysis metabolism in the blastocyst, enhanced TCA cycle activity from the 8-cell embryo stage, and active glycolysis in the blastocyst. Besides the metabolites detected at high levels in the zygote, which may be derived from oogenesis, most metabolites tend to increase during the later stages, consistent with the high energy demands associated with the transition of early embryos from a metabolically quiescent state to an active state. Together, these dynamic changes in metabolic signatures can be used to predict embryo quality and improve *in vitro* culture systems for assisted reproduction.

NAD^+^ is a prominent redox cofactor and enzyme substrate that is essential for energy metabolism, DNA repair, and epigenetic homeostasis. NAD^+^ repletion has been reported to restore oocyte quality, and enhances the ovulation rate and fertility in aged mice.^35^ However, the effect of NAD^+^ on mammalian pre-implantation development, and its mechanism, have remained elusive. In this study, we demonstrated that the depletion of NAD^+^ biosynthesis from the NAM salvage pathway, and the transition of pyruvic acid to lactate, but not the kynurenine pathway, resulted in mouse embryo developmental arrest at the cleavage stage, with the expression of many minor ZGA genes prolonged to late 2- and 4-cell stages. The results were in line with the notion that to maintain NAD^+^ levels, most NAD^+^ is recycled via salvage pathways rather than generated *de novo*. However, interestingly, we found without NAD^+^ from the NAM salvage pathway, the embryo development was not affected outside of this temporal window from 2-cell to 4-cell stage. We purpose that the normal development of FK866-treated embryos from zygote to 2-cell stage may depend on the large amount of maternal NAD^+^. Whereas, as the demonstration in the previous study that during the development the metabolic plasticity is increased, defined as flexibility in energetic substrate choice^16^, after the 4-cell stage, there may be other pathways to compensate for the intra-embryonic NAD^+^. For example, nicotinic acid can be converted to nicotinic acid mononucleotide by nicotinic acid phosphoribosyltransferase. Thus, the great need of NAD^+^ from 2- and 4-cell stages highlights the important role of the NAM salvage pathway. Notably, *Nampt* homozygous knockout (*Nampt*^−/−^) resulted in lethality within E5-10 days^36^, but not the 4- to 8-cell stages shown in our study. That may be due to the existence of maternal NAMPT, indicating the importance of maternal factors for embryonic development. Genome activation in the pre-implantation embryo occurs in two phases: minor and major ZGA. Major ZGA and the reprogramming of gene expression that occurs during major ZGA are critical for further development beyond the 2-cell stage, and minor ZGA, which is characterized by promiscuous transcription, supports major ZGA and is required for early embryo development.^37^ The transcription characteristics of minor ZGA observed in the late 2-cell embryo disrupted further embryo development. For example, the prolonged expression of *dux* significantly induced developmental arrest.^38^ Thus, developmental deficiency induced by the depletion of NAD^+^ can at least partly be attributed to excessive minor ZGA.

Although NAD^+^ is often used as a cofactor or substrate across a range of reactions, we found that excessive minor ZGA and developmental arrest induced by NAD^+^ depletion was mainly contributed by the failure of zyH3K27ac erasure, and were recapitulated by knockdown of the NAD^+^-dependent deacylase SIRT1. Therefore, we concluded that NAD^+^, in cooperation with SIRT1, plays an important role in the removal of zygotic H3K27ac. H3K27ac is a key acetylation histone that loosens the chromatin state and increases the accessibility of RNA polymerase II, resulting in gene expression.^30,39^ Chromatin may exist in a markedly relaxed state after fertilization, followed by progressive maturation of higher-order chromatin architecture during early development.^40^ Similarly, high enrichment of H3K27ac, which is associated with the loosened chromatin state, was observed in the zygote and early 2-cell embryo, and its removal resulted in a tighter state in late 2- and 4-cell embryos. Pol II reloading has previously been demonstrated in the zygote; in this study, we demonstrated that the reloading at the promoters of minor ZGA genes with the loosened chromatin state, associated with the zyH3K27ac enrichment, facilitated the minor ZGA. Thus, failed zyH3K27ac erasure may result in the maintenance of Pol II reloading and induce the prolonged expression of minor ZGA genes. In addition, pyruvic acid-dependent histone acetylation has been linked to ZGA. However, pyruvic acid -deprived embryos did not show a reduction in H3K27ac at the zygote stage (Supplementary Fig. S5o). Therefore, we propose that the development block caused by pyruvic acid deprivation may be related to a loss of NAD^+^ produced by the transition of pyruvic acid to lactate, suggesting that large amounts of NAD^+^ are needed by the 2-cell embryo. Most strikingly, in human embryos, we also detected the removal of zyH3K27ac at 8-cell stage that precisely matches the timing of major ZGA in human, and the NAD^+^ supplement could promote the removal and benefit the preimplantation development. Thus, the mechanism of NAD^+^-SIRT1-mediated deacetylation seems to be conserved among mammals. It is worth noting that the improvement of human embryonic development by NMN addition may be due to the employment of the aged MII oocytes from the *in vitro* maturation of immature human GV-stage oocytes obtained during normal human superovulation cycles in this study. Postovulatory aging is known to compromise the oocyte quality as well as subsequent embryo development in many different animal models, and becomes one of the most intractable issues that limit the outcome of human assisted reproductive technology (ART).^41,42^ In mice, it has been proved that NMN could improve the embryonic developmental potential of aged mouse oocytes.^35^ Thus, we purpose that NMN may be effective in improving the developmental potential of IVM oocytes in human.

In summary, we explored the characteristic metabolic patterns of mouse pre-implantation embryo development. Our dataset represents a framework for future studies to elucidate the complex interplay between metabolism and embryo development, and will provide tremendous opportunities to define better culture systems for human-assisted reproduction. We also found that the use of NMN as an NAD^+^ precursor promoted mammalian pre-implantation embryo development. Future studies should test NMN and other NAD^+^-increasing compounds in the clinical setting, as an additive to embryo medium, to improve human embryo quality and viability. Although these results are promising, we caution against the use of NAD^+^-increasing supplements until clinical studies have been completed.

## MATERIALS AND METHODS

### Experimental model and subject details

#### Ethics statement

This study was approved by the Institutional Review Board (IRB) of Chongqing Health Center for Women and Children (2019-603), China. In accordance with the measures of the People’s Republic of China on the administration of Human Assisted Reproductive Technology, the ethical principles of the Human Assisted Reproductive Technology and the Human Sperm Bank as well as the Helsinki declaration. The research followed the guiding principles of the Human Embryonic Stem Cell Ethics issued by the MOST and MOH and was regularly reviewed by the Medical Ethics Committee of Chongqing Health Center for Women and Children.

#### Human gamete and early embryo collection

For sperm, SpermGrad (Vitrolife) was used for gradient sperm separation. For oocytes, the controlled ovarian stimulation was carried out using GnRH analogues combined with human menopausal gonadotrophins or recombinant follicle stimulating hormone (FSH) for pituitary desensitization. Transvaginal ultrasound-guided oocyte collection was scheduled 36 h after hCG administration. All MII oocytes were selected for Intra cytoplasmic sperm injection (ICSI), while those identified as immature (germinal vesicle (GV) oocytes) were kept in incubator for 4–6 h. These GV oocytes matured *in vitro* to MII oocytes and could be used as a back-up reserve for clinical purpose. Once fertilization and formation of good embryos were achieved, the remaining *in vitro* maturation (IVM) MII oocytes superfluous to clinical need were donated to this study. These oocytes were obtained from women aged 25-37 years old. Each batch of embryos was equally divided into control and NMN-treated groups. ICSI was used in this study. Fertilization was assessed 17-20 h after ICSI. Numbers of embryos used for this study are: zygote (2PN): 22; 2-cell embryos: 2; 8-cell embryos: 2. The human embryos were cultured in Vitrolife sequential media system G1/G2 culture solution at 37°C, 6% CO_2_ and 5% O_2_. For NMN supplement, the 2PN zygotes were cultured in G1 media in the presence of NMN (10 µM, 1N3501; Sigma). All human embryos were cultured in G2 medium in the absence of NMN.

#### Mice

All animals used in this study were maintained and handled according to policies approved by Chongqing Health Center for Women and Children. Male and female ICR mice (6–8 weeks old) were purchased from Charles River (Beijing, China).

#### Embryo collection

Embryos were collected from superovulated female ICR mice treated with 6.5 IU of pregnant mare serum gonadotropin (PMSG); after 45-47 h, they were treated with 5 IU of human chorionic gonadotropin (hCG) and crossed to ICR males. Embryos were collected at the following times post-hCG injection: 24 h for the zygote stage, 34 h for the early 2-cell stage, 41 h for the middle 2-cell stage, 48 h for the late 2-cell stage, 58 h for the 4-cell stage, 69 h for the 8-cell stage, and 94 h for the blastocyst stage. Additionally, zygotes were collected at the following times: 12 h for the PN0 stage, 15 h for PN1, 18 h for PN2, 21 h for PN3, 24 h for PN4, and 27 h for PN5. Embryos were cultured in KSOM medium (MR-121-D; Merck, Kenilworth, NJ, USA) under mineral oil at 37°C in a 5% CO_2_ incubator.

#### Treatment of oocytes with inhibitors and chemicals

For inhibitor treatment, we prepared solutions of FK866 (HY-50876; MedChemExpress, Monmouth Junction, NJ, USA), GEN-140 (HY-100742A; MedChemExpress), Oxamate (C3893; ApexBio, Houston, TX, USA) and TSA (T8552; Sigma, St. Louis, MO, USA) in dimethyl sulfoxide (DMSO). NMN (N3501; Sigma) solutions were prepared in water. Solutions were diluted to yield a final concentration in maturation medium as needed (FK866, 0.01 µM; GNE-140, 20 µM; Oxamate, 20 mM; NMN, 10 µM; TSA, 10 nM). Embryos were cultured *in vitro* in KSOM medium containing different doses of inhibitor for further analysis. Correspondingly, 0.05% DMSO was included as a negative control. To assess the effects of NAD^+^ synthesis on early embryonic development, zygotes cultured in KSOM medium were supplemented with different concentrations of FK866, GNE-140 and Oxamate. The relevant phenotypes were examined at the indicated time points.

#### Metabolomics

Mouse embryos were washed in PBS and snap-frozen in liquid nitrogen for 15 min, and were stored at −80 °C in cold 80% methanol. 200 μL of extraction solution buffer (methanol: acetonitrile: water = 2:2:1 (V/V)) was added to the sample, freeze in liquid nitrogen for 1 min, thaw and vortex for 30s. Repeat the above operation. Sonicate the samples in an ice-water bath for 10 min, and incubate at -40 °C for 1 h. Centrifuge the samples at 4 °C for 15 min at 12,000 × rpm, and the supernatant was transferred to a new tube (pre-chilled on dry ice) and evaporated with a speed vacuum. Dried metabolites were reconstituted in 30 μl of 0.03% formic acid in water, vortexed, centrifuged at 14,000g for 15 min at 4 °C. All samples were mixed into QC samples with the same amount of supernatant, and the supernatant was analysed using an UHPLC device (1290 Infinity LC; Agilent Technologies, Santa Clara, CA, USA) coupled to a quadrupole time-of-flight system (AB Sciex TripleTOF 6600; Shanghai Applied Protein Technology Co., Ltd., Shanghai, China). For hydrophilic interaction chromatography (HILIC) separation, samples were analyzed using a 2.1 mm × 100 mm Acquity UPLC BEH 1.7 µm column (Waters, Milford, MA, USA). In both electrospray ionization (ESI)-positive and -negative modes, the mobile phase contained A = 25 mM ammonium acetate and 25 mM ammonium hydroxide in water and B = acetonitrile. The gradient was 85% B for 1 min; this was linearly reduced to 65% in 11 min, reduced to 40% in 0.1 min and held for 4 min, and then increased to 85% in 0.1 min, with a 5 min re-equilibration period. The ESI source conditions were set as follows: ion source gas 1 (Gas 1), 60; ion source gas 2 (Gas 2), 60; curtain gas (CUR), 30; source temperature, 600°C, ion spray voltage floating (ISVF), 5,500 V. In MS-only acquisition, the instrument was set to acquire over an m/z range of 60–1,000 Da, and the accumulation time for TOF-MS scanning was set at 0.20 s/spectra. In auto tandem mass spectrometry (MS/MS) acquisition mode, the instrument was set to acquire over an m/z range of 25–1,000 Da, and the accumulation time for product ion scanning was set at 0.05 s/spectra. We used the Collection of Algorithms of MEtabolite pRofile Annotation (CAMERA) to annotate isotopes and adducts. Only variables having > 50% non-zero measurement values in at least one group were extracted. Metabolite compounds were identified by comparing m/z values (< 25 ppm) and MS/MS spectra with an in-house database established using available authentic standards. Chromatogram review and peak area integration were performed using MultiQuant software v.3.0 (SCIEX). Finally, 246 metabolites of common and important metabolic pathways (for example, energy metabolism, carbohydrate metabolism, amino acid metabolism and nucleotide metabolism) were included in this method. Each condition was assayed in sextuplicate, with each replicate containing approximately 1,000 embryos, the exact number of which was used to normalize the data. For drawing the Sankey diagram, the highest and lowest values were determined by normalizing the levels of metabolites in each stage of the early embryos, and the tertiles were defined as high, intermediate and low levels, respectively.

#### In vitro transcription of Trim21 mRNA

To prepare mRNAs for microinjection, pGEMHE-mCherry-mTrim21 (105522; Addgene, Watertown, MA, USA) vectors were linearized and purified with phenol-chloroform, followed by ethanol precipitation. Linearized DNAs were transcribed *in vitro* using the mMESSAGE mMACHINE Kit (AM1344; Invitrogen, Carlsbad, CA, USA). Transcribed mRNAs were added to poly (A) tails (200–250 bp) using the mMACHINE Kit (AM1350; Invitrogen), recovered by lithium chloride precipitation, and resuspended in nuclease-free water. *Trim21* mRNA was frozen and stored at −80°C. For trimming, *Trim21* mRNA (1 µg/µL), *Sirt1* antibodies (0.5 µg/µL), and siRNA (10 µM) were mixed and co-injected into zygotes through microinjection.

#### Microinjection

All siRNAs used in this study were purchased from RiboBio. The scrambled siRNA and *Trim21* mRNA was injected into the zygotes as the negative control. We placed zygote-stage embryos in 150 µg/mL hyaluronidase (4272; Sigma) to digest the outer granule cells. The siRNA was centrifuged at 12,000 rpm for 10 min at 4°C, and stored at 4°C until use. Then, microinjection was performed using a FemtoJet 4i microinjector (Eppendorf, Hamburg, Germany) and E LIPSE Ti micromanipulators (Nikon, Tokyo, Japan). For injection, a glass capillary Femtotip (Eppendorf) was loaded with 2 μL of RNA mixtures using a Microloader (Eppendorf), and the solution was injected into the cytoplasm in a drop of M2 medium (M7167; Merck). The injection volume was approximately 2–5 pL. The injection conditions consisted of an injection pressure of 250 hPa, compensation pressure of 60 hPa, and injection time of 0.7 s. Immediately after microinjection, embryos were cultured in KSOM medium at 37°C in 5% CO_2_.

#### Immunofluorescence staining

After removing the zona pellucida with acidic operating fluid, mouse embryos were fixed in fixative solution (FB002; Invitrogen) for 40 min at room temperature, followed by permeabilization in 1% Triton X-100 (93443, 100 ml; Sigma) for 20 min at room temperature. Embryos were then blocked in blocking solution consisting of 1% bovine serum albumin (BSA) in phosphate-buffered saline (PBS) for 1 h at room temperature after three washes in washing solution (0.1% Tween-20, 0.01% Triton X-100 in PBS). Antibody incubation (DUX: sc-137190; Santa Cruz Biotechnology, Santa Cruz, CA, USA; SIRT1: 8469S; CST, Danvers, MA, USA; H3K27ac: 39034; Active Motif, Carlsbad, CA, USA) was performed overnight at 4°C. The next day, the embryos were washed in washing solution and incubated with secondary antibodies (A10040; Invitrogen) for 1 h at room temperature. After staining with Hoechst, the embryos were washed in washing solution. Embryo imaging was performed using an inverted confocal microscope (TCS SP8; Leica, Wetzlar, Germany) and analyzed using LAS X software (Leica).

#### qPCR

Total RNA was extracted using the Arcturus PicoPure RNA Isolation Kit (12204-01; Ambion, Austin, TX, USA) according to the manufacturer’s instructions, and reverse transcription was performed to generate cDNA using the PrimeScript RT Master Mix (RR036; Takara, Kusatsu, Japan). The RNase-Free DNase Set (79254; Qiagen, Hilden, Germany) was used to ensure that there was no DNA contamination. We performed qPCR using the TB Green Premix Ex Taq II (RR820; Takara) and CFX96 Real-Time System (Bio-Rad, Hercules, CA, USA). The reaction parameters were as follows: 95°C for 30 s, followed by 40 two-step cycles of 95°C for 5 s and 60°C for 34 s. Hprt was used as a reference gene. Ct values were calculated using the CFX96 system software, and the target sequences were normalized to the reference sequence using the 2−^ΔΔCt^ method.

#### Measurement of NAD^+^ and NADH levels

NAD^+^ and NADH levels were determined using the NAD^+^/NADH Glo cycling assay (G9071; Promega, Madison, WI, USA). Embryos were lysed with 150 µL 0.2 M NaOH and incubated for 10 min at room temperature. The sample solution was divided into two tubes (50 µL/tube). For NAD^+^ measurement, 25 µL 0.4N HCl was added to the solution, which was heated at 60°C for 15 min. After 10 min of incubation at room temperature, 25 µL 0.5 M Trizma base (T1503; Sigma) was added to the resulting samples. For NADH measurement, the sample solution was heated at 60°C for 15 min and incubated for 10 min at room temperature, followed by the addition of 25 µL 0.5M Trizma base and 25 µL 0.4N HCl. The assay reagent was added to the sample solution, which was incubated for 3 h prior to cycling; luminescence was then determined (Varioskan LUX; Thermo Fisher, Waltham, MA, USA).

#### RNA-seq

RNA-seq libraries were prepared as previously described. Briefly, three embryos were used per reaction, and two replicates were performed for each group. All embryos were washed three times in 0.5% BSA-PBS solution to avoid possible contamination. cDNA was amplified using the Phusion Hot Start II High-Fidelity PCR Master Mix (F-565S; Thermo Fisher). Library preparation was performed using the NEBNext Ultra II DNA Library Prep Kit (E7805S, New England Biolabs, Ipswich, MA, USA) according to the manufacturer’s instructions. All libraries were sequenced using the NovaSeq 6000 (Illumina, San Diego, CA, USA) according to the manufacturer’s instructions.

#### RNA-seq data processing

For RNA-seq analysis of early stage embryos, FastQC was performed for Illumina reads. We used the Trim Galore software to discard low-quality reads, trim adaptor sequences, and eliminate poor-quality bases. Then, we downloaded the mouse reference genome (genome assembly: mm10) from the Ensembl database and used the HISAT2 software for read alignment. The gene-level quantification approach was used to aggregate raw counts of mapped reads using the featureCounts tool. The expression level of each gene was quantified in terms of the normalized fragments per kilobase of transcript per million mapped reads (FPKM). Next, we used the R package *DESeq2* for differential gene expression analysis. KEGG analysis of screened DEGs was performed using the KOBAS online tool (http://kobas.cbi.pku.edu.cn/kobas3/).

#### CUT&Tag

CUT&Tag was performed using the Hyperactive In-Situ ChIP Library Prep Kit for Illumina (TD901; Vazyme Biotech, Nanjing, China). Embryos were incubated with 10 μl pre-washed ConA beads. 50 μL of antibody buffer; 0.5 μg antibody was added and the mixture was cultured for 1.5 h at room temperature. After washing twice with dig-wash buffer, 150 μL dig-wash buffer with 0.7 μg secondary antibody was added and the mixture was incubated at room temperature for 1 h. After washing twice with 800 μL dig-wash buffer, 0.3 μL pG–Tn5 and 50 μL dig-300 buffer were added, and the samples were incubated at room temperature for 1 h and then washed twice with 800 μL dig-wash buffer. We added 300 μL of tagmentation buffer and the samples were incubated at 37°C for 1 h. The reaction was stopped with 10 μL 0.5 M ethylenediaminetetraacetic acid (EDTA), 3 μL 10% sodium dodecyl sulfate (SDS) and 2.5 μL 20 mg/ml proteinase K. After extraction with phenol-chloroform and ethanol precipitation, PCR was performed to amplify the libraries under the following cycling conditions: 72°C for 5 min, 98°C for 30 s, 20 cycles of 98°C for 10 s, and 60°C for 30 s, followed by a final extension at 72°C for 1 min and holding at 4°C. Post-PCR clean-up was performed by adding a 1.5× volume of DNA Clean Beads (N411; Vazyme Biotech), and libraries were incubated with beads for 10 min at room temperature, washed gently in 80% ethanol, and eluted in 20 μL water. All libraries were sequenced using the Illumina NovaSeq 6000 platform according to the manufacturer’s instructions.

#### CUT&Tag data analysis

We aligned paired-end CUT&Tag reads of H3K27ac to the mm10 reference genome using Bowtie2 v2.2.2 software, with the following options: -local -very-sensitive -local-no-unal -nomixed -no-discordant -phred33 -I 10 -X 700, followed by filtering with SAMtools v1.7 software at MAPQ5. Unmapped and non-uniquely mapped reads were removed. CUT&Tag data reproducibility between replicates was assessed by correlation analysis of mapped read counts across the genome. Then, we pooled the biological replicates for each stage and performed downstream analysis. For peak calling, we converted sequence alignments in BAM format into BEDPE records using the BEDTools function *bamtobed*. CUT&Tag peaks were called on merged replicates and normalized to input using the MACS2 v2.1.1.20160309 software. Peak calls with a false-discovery rate (FDR) of ≤ 5% were used for downstream analysis. Peak annotation was performed using the R package *ChIPseeker* with the default parameters. Peak comparisons and overlaps were evaluated using the BEDTools suite for autosomal chromosomes. For quantitative analysis, we normalized the read counts by computing the number of reads per kilobase of transcript per million mapped reads (RPKM). RPKM values were calculated using merged replicate BAM files using the *bamCoverage* tool in deepTools software. To minimize batch and cell type variation, the RPKM values were further normalized through Z-score transformation (whole-genome 100-bp bins, excluding outliner regions). The washU epigenome browser (http://epigenomegateway.wustl.edu/browser/) was used to visualize H3K27ac CUT&Tag data.

#### Identification of aberrant H3K27ac domains

For the late 2- and 4-cell stages, we quantified the expression levels for each putative H3K27ac domain using the BEDTools coverage tool, and then compared H3K27ac levels between the control and treatment groups using the *edgeR* package in R. The aberrant H3K27ac domains of each stage were identified using stringent criteria: log2(fold change) > 1.5 or < −1.5, and sum of the normalized H3K27ac levels of the control and treated groups ≥ 2. The retained regions were merged using the *merge* function in BEDTools v2.27.1 if they were consecutive or overlapped.

#### ZGA genes within H3K27ac domains

The predefined ZGA genes list was from the previous study.^25^ Promoters of the ZGA genes (defined as ±1 kb around the transcription start site) that overlapped with H3K27ac peaks according to the BEDTools *intersect* tool were defined as ZGA gene H3K27ac domains. Changes in CUT&Tag data were processed and visualized using heatmaps or metaplots produced using the *computeMatrix* and *plotHeatmap* functions of deepTools. Data were imported using the default settings and all values were normalized to RPKM values and scaled as described above.

#### ATAC-seq data processing and analysis

ATAC-seq data were analyzed using a standardized software pipeline developed by the ENCODE DCC for the ENCODE Consortium to perform quality-control analysis and reads alignment. ATAC–seq reads were trimmed with a custom adaptor script and mapped to mm10 using Bowtie2 v2.2.2 and Samtools v1.7 to eliminate PCR duplicates and filter alignment reads with mapping quality scores (MAPQ) > 30. To analyze the ZGA gene, we produced bigWig coverage profiles from BAM files by deepTools v3.5.1. And the 1 kilobases upstream and downstream scores of per ZGA gene promoter was calculated and visualized using computeMatrix and plotProfile from deepTools.

#### Pol II ChIP-seq data processing and analysis

The paired-end Pol II ChIP-seq reads were aligned with the parameters: -t -q -N 1 -L 25 -X 2000-no-mixed-no-discordant by Bowtie v2.2.2. All unmapped reads, non-uniquely mapped reads and PCR duplicates were removed by Picard v2.25.7. The ZGA gene promoter Pol II enrichment was calculated through the scores of 1 kilobases upstream and downstream of gene boundary.

#### Statistical analyses

All statistical analyses were performed using R v4.1.0 software (R Development Core Team, Vienna, Austria). Data are expressed as means ± standard error of the mean (SEM). Differences between means were evaluated using the two-tailed Student’s *t*-test or Wilcoxon rank sum test. Asterisks indicate significant differences as follows: *P < 0.05, **P < 0.01, and ***P < 0.001.

## Acknowledgements

We thank Xiaoyu Hu and Wei Xie from Tsinghua University for their technical assistance, and the generous donors whose contributions have enabled this research. This work was supported by the Basic public welfare research program of Zhejiang Province (LY22C120001), Chongqing Science and Health Joint Project (2021MSXM072); Special Research Project of Chongqing Key Laboratory of Human Embryo Engineering, National Natural Science Foundation of China (82070981).

## Author Contributions

J.L., J.Z., S.G., G.H. and Q.K. conceived and designed the study. J.Z. and X.Y. performed mouse embryo collection. J.L. performed human embryo experiments with contributions from G.H., Z.L., M.G. and M.Z. J.Z. performed all mouse embryo experiments with contributions from Y.Z., X.L. and Z.D. W.H. and J.Z. analyzed the data with contributions from Q.K., X.Y., J.S., W.C. and C.D. S.G., G.H. and Q.Z. supervised the project. J.Z., S.G. and Q.K. wrote the manuscript.

## Notes

### Competing Interest Statement

The authors have declared no competing interest.

