## Supplemental Information for "Metabolic control of histone acetylation for precise and timely regulation of minor ZGA in early mammalian embryos"

Supplementary Information:

Supplementary Figure S1-S7

Supplementary Table S1-S6

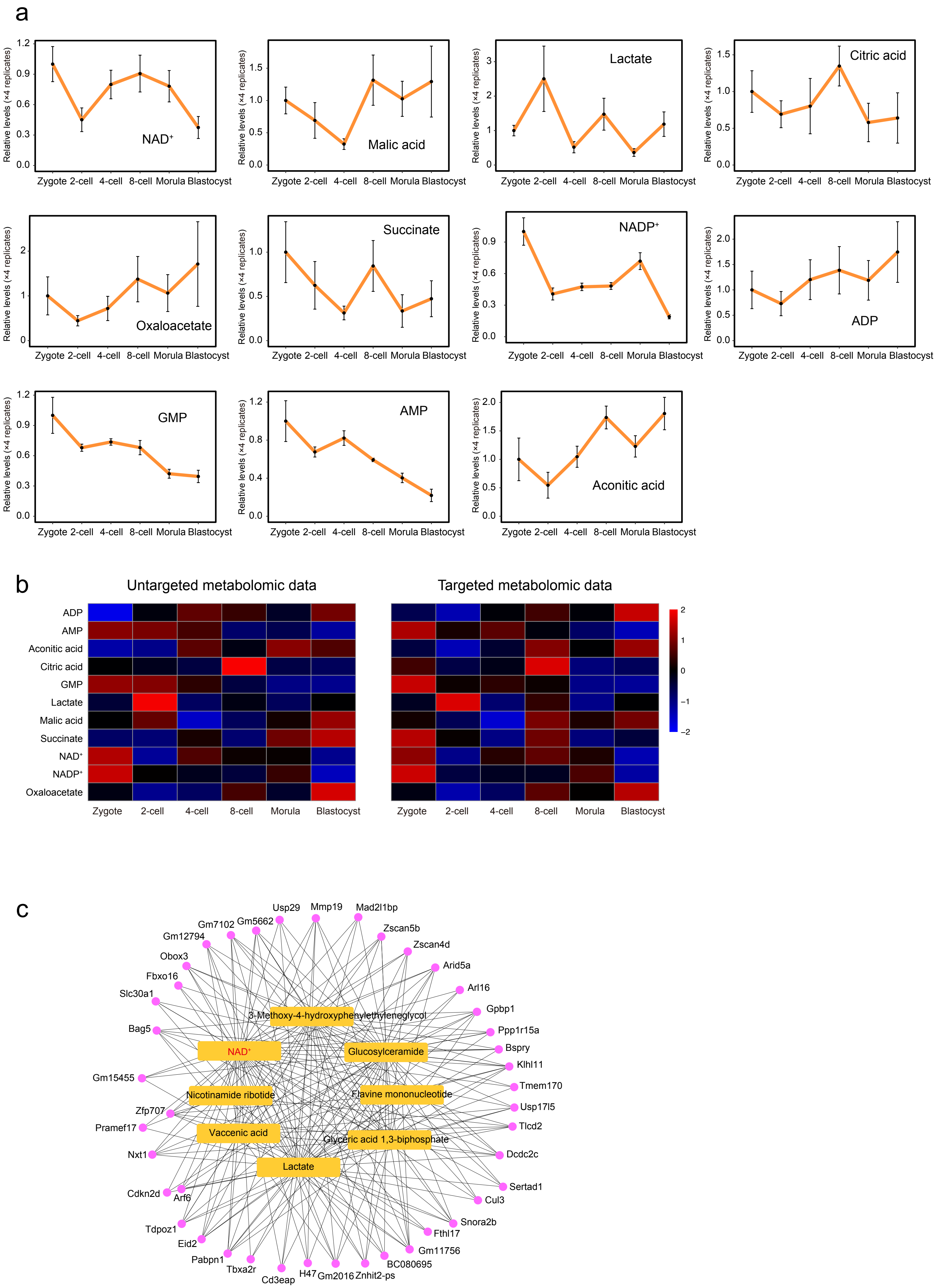
a
Relative levels (×4 replicates)
Zygote 2-cell 4-cell 8-cell Morula Blastocyst
NAD⁺
Malic acid
Lactate
Citric acid
Oxaloacetate
Succinate
NADP⁺
ADP
GMP
AMP
Aconitic acid
b
Untargeted metabolomic data
Targeted metabolomic data
ADP
AMP
Aconitic acid
Citric acid
GMP
Lactate
Malic acid
Succinate
NAD⁺
NADP⁺
Oxaloacetate
Zygote
2-cell
4-cell
8-cell
Morula
Blastocyst
2
1
0
−1
−2
c
3-Methoxy-4-hydroxyphenylethyleneglycol
NAD⁺
Glucosylceramide
Nicotinamide ribotide
Flavine mononucleotide
Vaccenic acid
Glyceric acid 1,3-biphosphate
Lactate
Usp29
Mmp19
Mad2l1bp
Zscan5b
Zscan4d
Arid5a
Arl16
Gpbp1
Ppp1r15a
Bspry
Klhl11
Tmem170
Usp17l5
Tlcd2
Dcdc2c
Sertad1
Cul3
Snora2b
Fthl17
Gm11756
BC080695
Znhit2-ps
Gm2016
H47
Cd3eap
Tbxa2r
Pabpn1
Eid2
Tdpoz1
Arf6
Cdkn2d
Nxt1
Pramef17
Zfp707
Gm15455
Bag5
Slc30a1
Fbxo16
Obox3
Gm12794
Gm7102
Gm5662

**Figure S1. Targeted metabolomic analysis. a** Relative level of 11 metabolites related to carbohydrate metabolism during mouse early embryo development based on targeted metabolomic data. **b** Heatmap showing comparison of untargeted and targeted metabolome data. **c** Network connections of metabolites (yellow) and genes (pink)

a

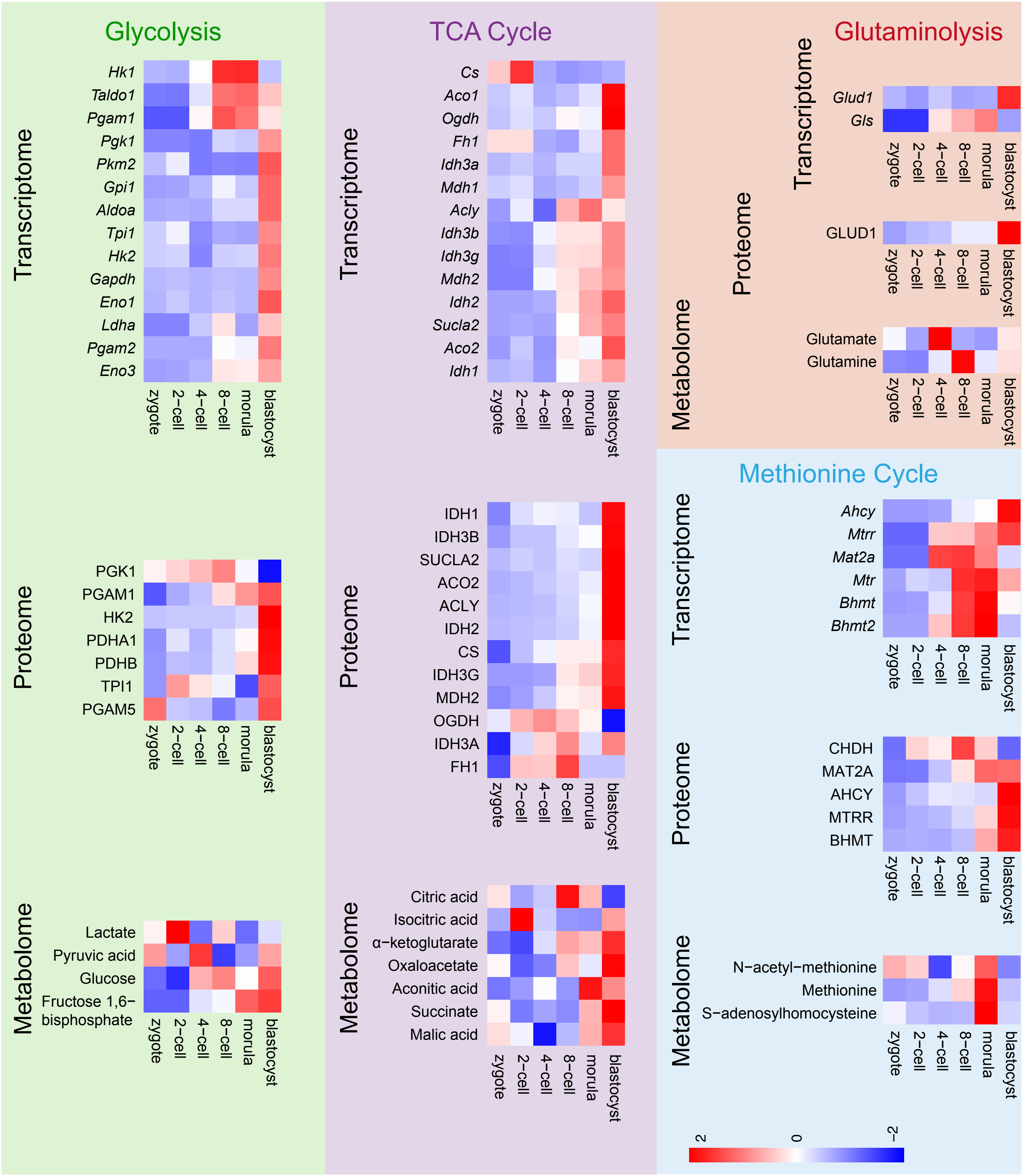

Glycolysis
Transcriptome
Hk1
Taldo1
Pgam1
Pgk1
Pkm2
Gpi1
Aldoa
Tpi1
Hk2
Gapdh
Eno1
Ldha
Pgam2
Eno3
zygote
2−cell
4−cell
8−cell
morula
blastocyst
Proteome
PGK1
PGAM1
HK2
PDHA1
PDHB
TPI1
PGAM5
Metabolome
Lactate
Pyruvic acid
Glucose
Fructose 1,6−bisphosphate
TCA Cycle
Cs
Aco1
Ogdh
Fh1
Idh3a
Mdh1
Acly
Idh3b
Idh3g
Mdh2
Idh2
Sucla2
Aco2
Idh1
IDH1
IDH3B
SUCLA2
ACO2
ACLY
IDH2
CS
IDH3G
MDH2
OGDH
IDH3A
FH1
Citric acid
Isocitric acid
α−ketoglutarate
Oxaloacetate
Aconitic acid
Succinate
Malic acid
Glutaminolysis
Glud1
Gls
GLUD1
Glutamate
Glutamine
Methionine Cycle
Ahcy
Mtrr
Mat2a
Mtr
Bhmt
Bhmt2
CHDH
MAT2A
AHCY
MTRR
BHMT
N−acetyl−methionine
Methionine
S−adenosylhomocysteine
2
0
−2

**Figure S2. Activities of metabolic pathways during mouse pre-implantation embryo development. a** Changes in metabolic enzyme levels during mouse pre-implantation development. Expression patterns of metabolic enzymes on RNA and protein according to transcriptomic and proteomic data on mouse early embryos.

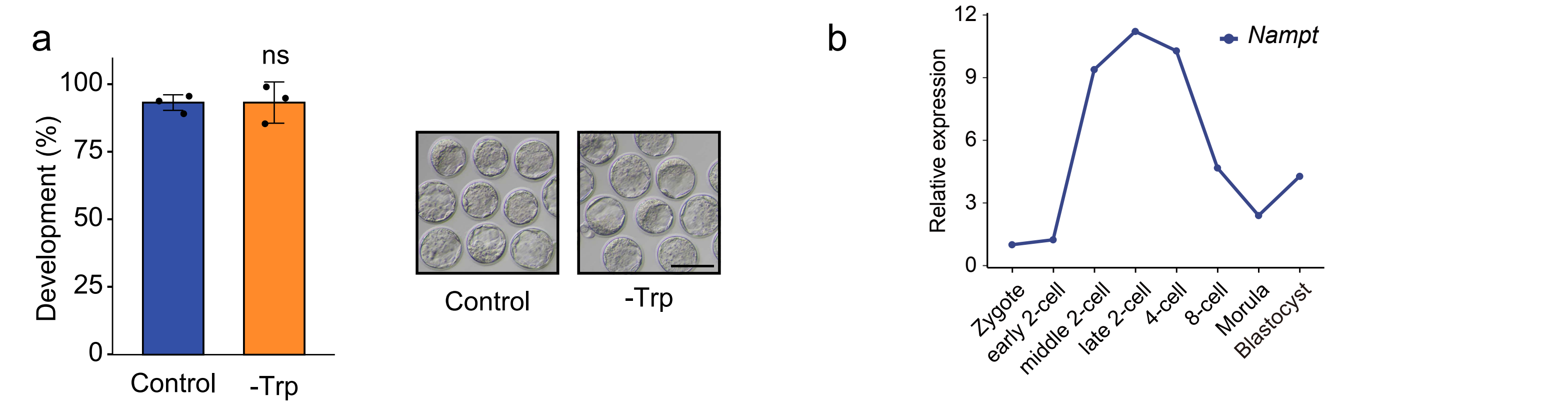


Expression level during preimplantation development (protein)

| Gene | Zygote | 2-cell | 4-cell | 8-cell | Morula | Blastocyst |
|---|---|---|---|---|---|---|
| NAMPT | 0.5781 | 0.7520 | 0.9063 | 1.0256 | 1.0508 | 1.6872 |

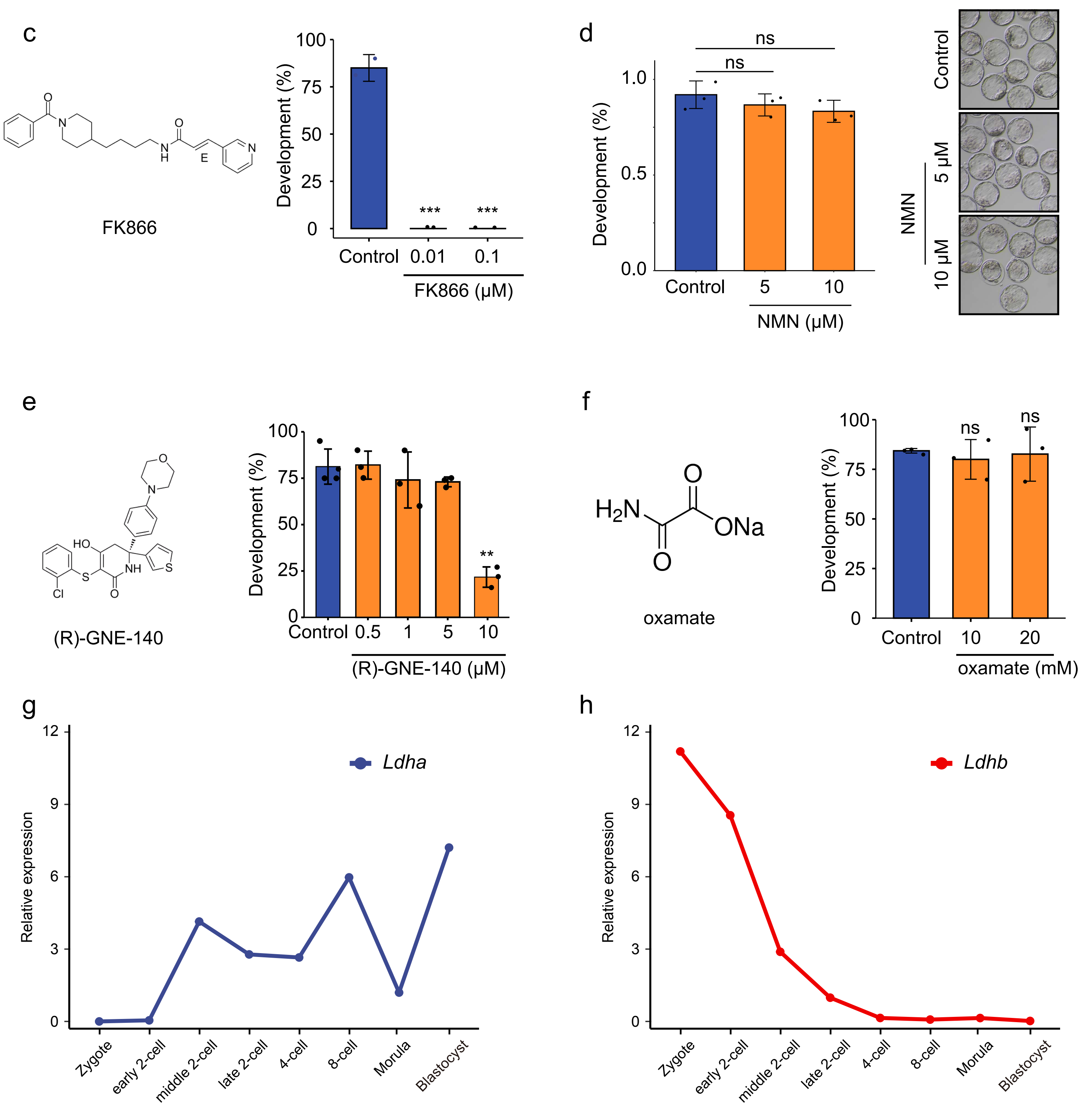

**Figure S3. Inhibition of NAD$^+$ synthesis impairs mouse early embryonic development.** **a**, Developmental rate of mouse embryos cultured in Trp-deprived medium. **b**, Expression patterns of *Nampt* at mRNA (upper) and protein (bottom) levels during mouse preimplantation embryonic development. **c**, Developmental rate of mouse embryos cultured in FK866-addition medium. **d**, Developmental rate of mouse embryos cultured in NMN-addition medium, 5 μM and 10 μM NMN were added into the KSOM medium. Neither concentrations could significantly affect the development of mouse embryos. Thus, the 10 μM NMN was used in the rescue experiments. Developmental rate of mouse embryos cultured in GNE-140-addition medium (**e**) and oxamate-addition medium (**f**). Expression patterns of *Ldha* (**g**) and *Ldhb* (**h**) according to RNA-seq data on mouse early embryos.

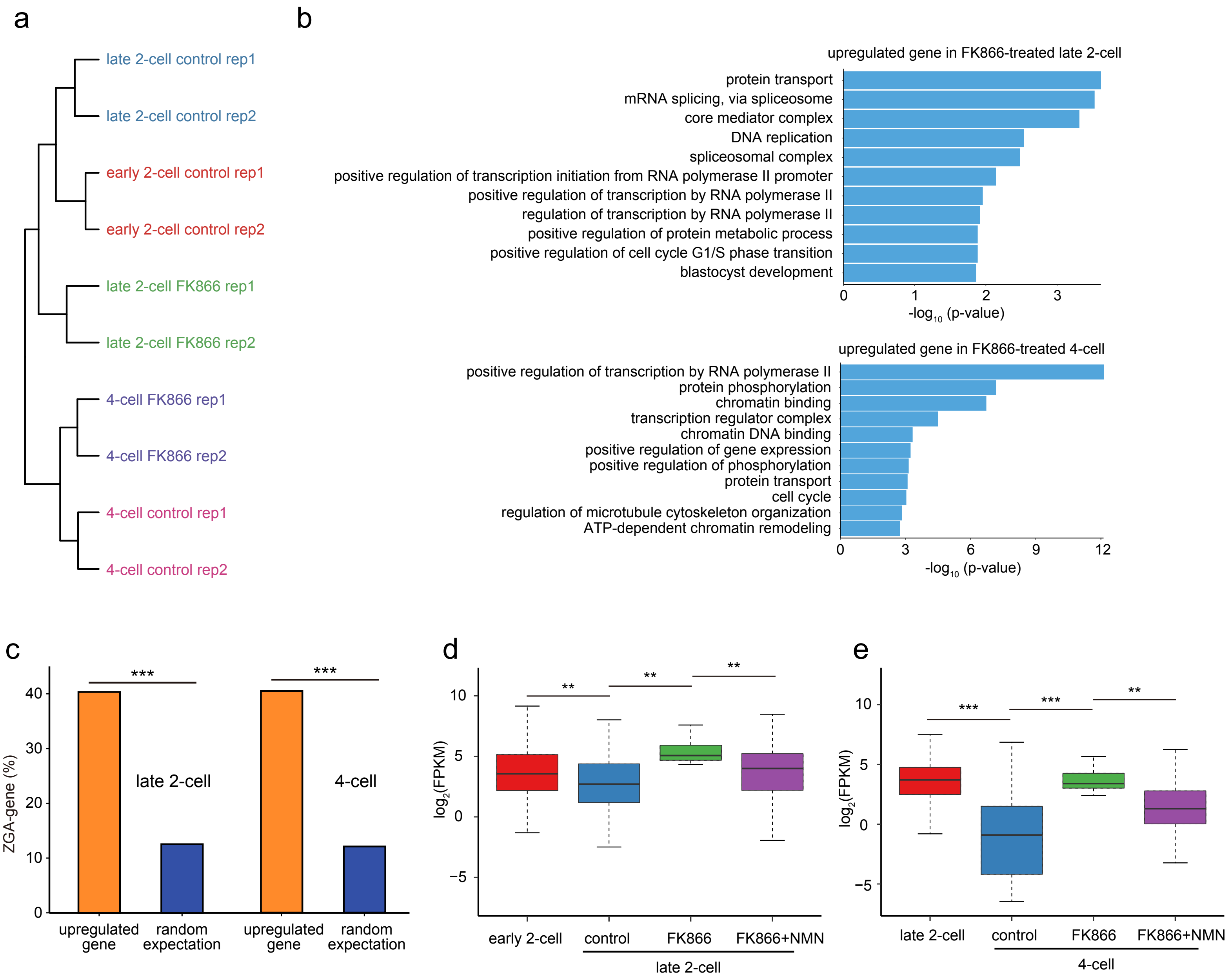

a
late 2-cell control rep1
late 2-cell control rep2
early 2-cell control rep1
early 2-cell control rep2
late 2-cell FK866 rep1
late 2-cell FK866 rep2
4-cell FK866 rep1
4-cell FK866 rep2
4-cell control rep1
4-cell control rep2
b
upregulated gene in FK866-treated late 2-cell
protein transport
mRNA splicing, via spliceosome
core mediator complex
DNA replication
spliceosomal complex
positive regulation of transcription initiation from RNA polymerase II promoter
positive regulation of transcription by RNA polymerase II
regulation of transcription by RNA polymerase II
positive regulation of protein metabolic process
positive regulation of cell cycle G1/S phase transition
blastocyst development
0 1 2 3
-log10 (p-value)
upregulated gene in FK866-treated 4-cell
positive regulation of transcription by RNA polymerase II
protein phosphorylation
chromatin binding
transcription regulator complex
chromatin DNA binding
positive regulation of gene expression
positive regulation of phosphorylation
protein transport
cell cycle
regulation of microtubule cytoskeleton organization
ATP-dependent chromatin remodeling
0 3 6 9 12
-log10 (p-value)
c
ZGA-gene (%)
0 10 20 30 40
***
***
late 2-cell
4-cell
upregulated gene
random expectation
upregulated gene
random expectation
d
log2(FPKM)
10 5 0 −5
** ** **
early 2-cell
control
FK866
FK866+NMN
late 2-cell
e
log2(FPKM)
10 5 0 −5
*** *** **
late 2-cell
control
FK866
FK866+NMN
4-cell

**Figure S4. RNA-seq analysis of control and FK866-treated mouse embryos. a** Unsupervised clustering of gene expression among control and FK866-treated embryos at the late 2- and 4-cell stages. **b** KEGG analysis of upregulated genes in FK866-treated late 2- and 4-cell embryos. **c** Bar plot showing the numbers of ZGA genes embedded within upregulated genes upon FK866 treatment, and those expected by chance in late 2- and 4-cell embryos. Box plots showing ZGA genes upregulated between control and FK866-treated embryos that were rescued by NMN supplementation in (**d**) late 2-cell and (**e**) 4-cell embryos, respectively.

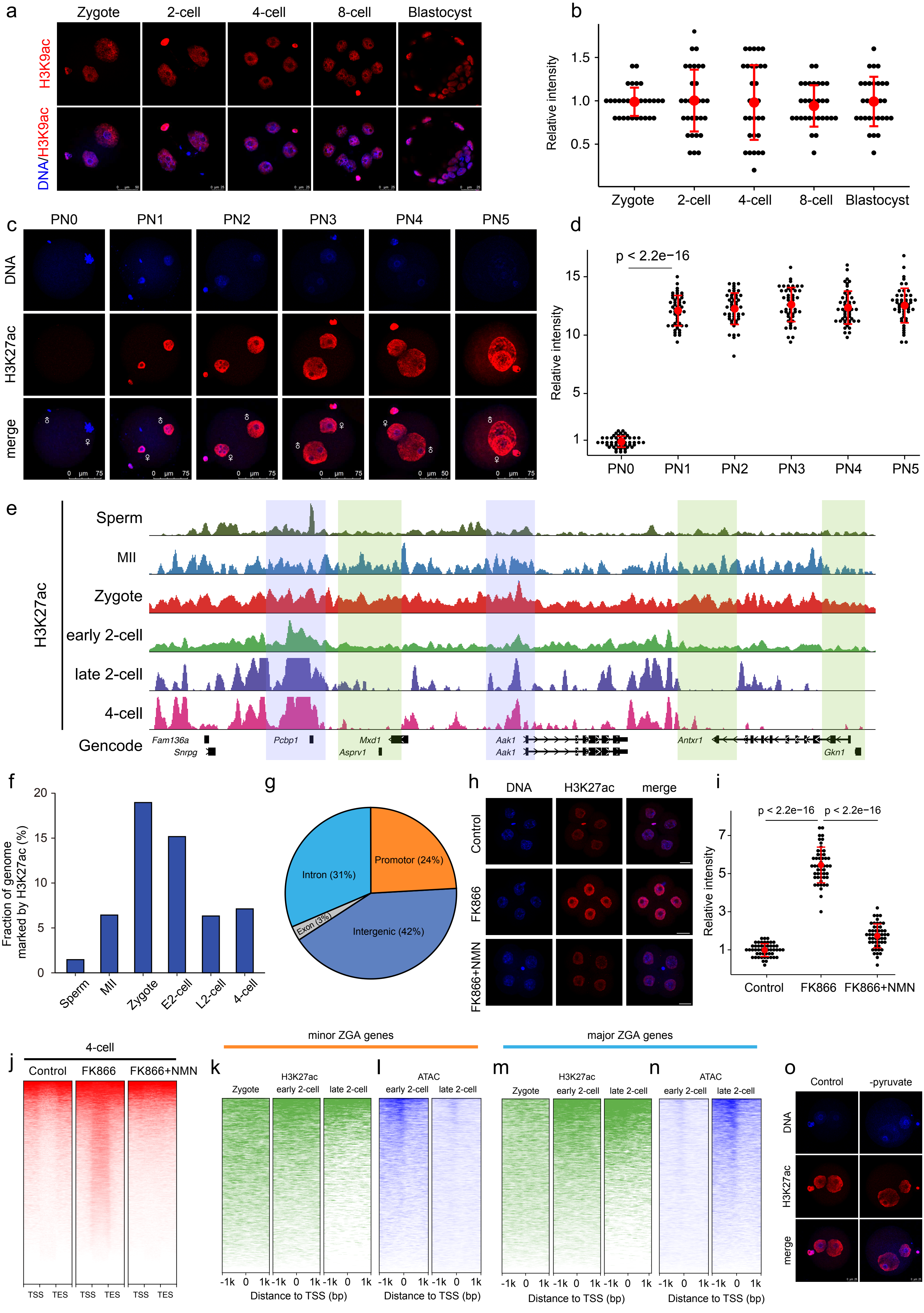

a
Zygote
2-cell
4-cell
8-cell
Blastocyst
H3K9ac
DNA/H3K9ac
b
Relative intensity
Zygote
2-cell
4-cell
8-cell
Blastocyst
c
PN0
PN1
PN2
PN3
PN4
PN5
DNA
H3K27ac
merge
d
p < 2.2e−16
Relative intensity
PN0
PN1
PN2
PN3
PN4
PN5
e
H3K27ac
Sperm
MII
Zygote
early 2-cell
late 2-cell
4-cell
Gencode
Fam136a
Snrpg
Pcbp1
Mxd1
Asprv1
Aak1
Aak1
Antxr1
Gkn1
f
Fraction of genome marked by H3K27ac (%)
Sperm
MII
Zygote
E2-cell
L2-cell
4-cell
g
Promotor (24%)
Intron (31%)
Exon (3%)
Intergenic (42%)
h
DNA
H3K27ac
merge
Control
FK866
FK866+NMN
i
p < 2.2e−16
p < 2.2e−16
Relative intensity
Control
FK866
FK866+NMN
j
4-cell
Control
FK866
FK866+NMN
TSS
TES
minor ZGA genes
major ZGA genes
k
H3K27ac
Zygote
early 2-cell
late 2-cell
l
ATAC
early 2-cell
late 2-cell
Distance to TSS (bp)
m
H3K27ac
Zygote
early 2-cell
late 2-cell
n
ATAC
early 2-cell
late 2-cell
Distance to TSS (bp)
o
Control
-pyruvate
DNA
H3K27ac
merge

**Figure S5. Establishment of zyH3K27ac landscapes in early mouse embryos. a, b** Immunostaining of H3K9ac during mouse pre-implantation embryo development. Representative image from three independent experiments is shown. Scale bar, 25 µm. Each dot represents a single nucleus. **c, d** Dynamics of H3K27ac enrichment across all pronuclear stages in the zygote. The H3K27ac fluorescence intensity of zygotes was quantified. Each dot represents a single nucleus. **e** Genome-wide landscapes of H3K27ac histone modification in mouse sperm, oocyte, zygote (PN4), early 2-cell, late 2-cell, and 4-cell embryos. **f** Fraction of the mouse genome covered by H3K27ac reads at different developmental stages. **g** Pie charts show the percentages of H3K27ac peaks assigned to the promoter, intron, exon, and intergenic regions. **h** Representative confocal images of control, FK866-treated, and FK866+NMN-treated 4-cell embryos stained with H3K27ac antibody. Scale bars, 25 µm. **i** Quantification of H3K27ac fluorescence intensity in 4-cell embryos. Each dot represents a single nucleus. **j** Heatmap showing all H3K27ac signals ranked by their relative change after FK866 treatment. NMN rescued the changes induced by FK866 treatment in 4-cell embryos. Heatmap showing minor ZGA genes (**k**) and major ZGA genes (**m**) promoter H3K27ac signals in zygote, early 2-cell, and late 2-cell embryos. Heatmap showing minor ZGA genes (**l**) and major ZGA genes (**n**) promoter ATAC signals in early and late 2-cell embryos. **o** Representative confocal images of control and pyruvate-deprived zygotes stained with H3K27ac antibody. Scale bars, 25 µm.

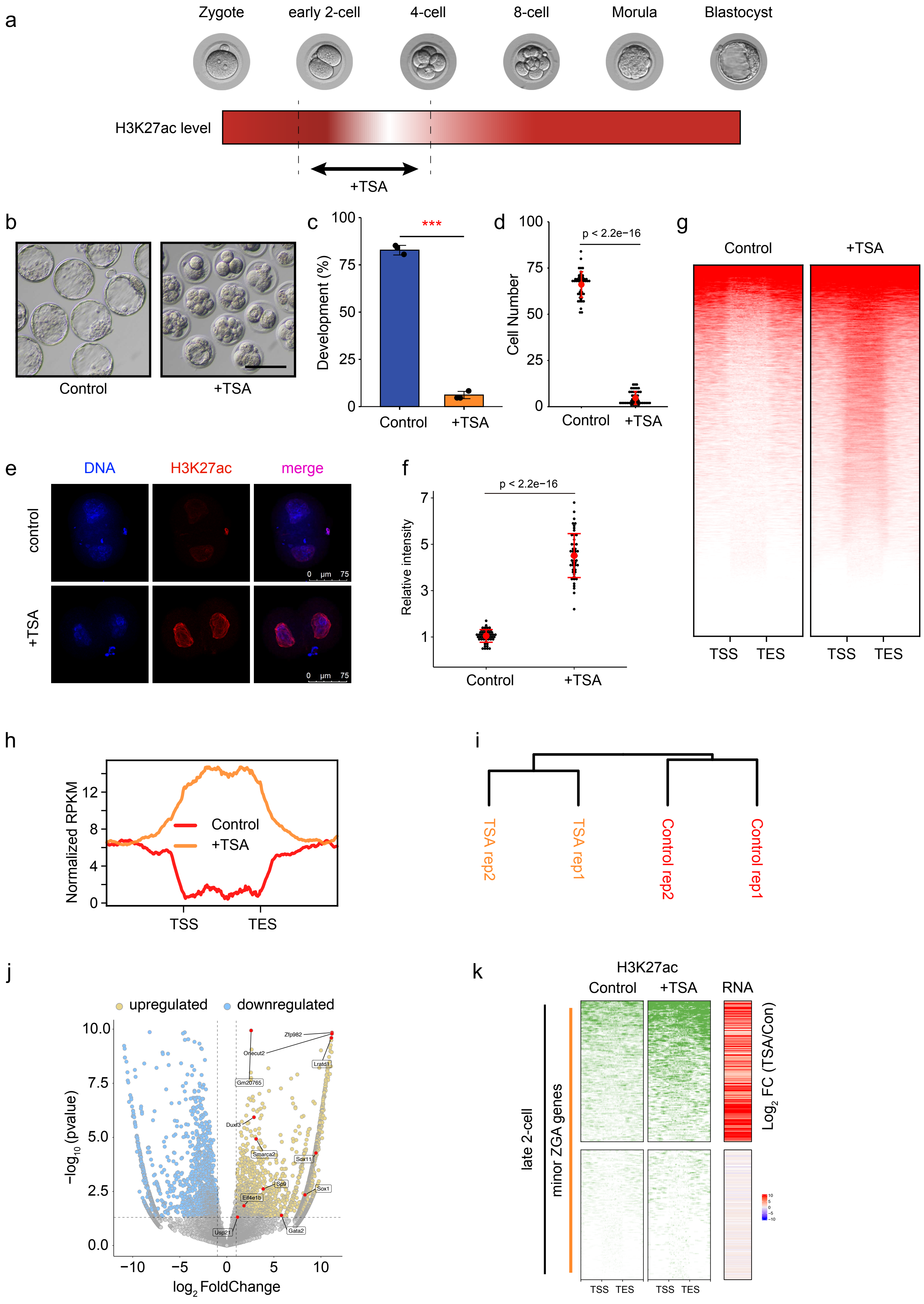

a
Zygote
early 2-cell
4-cell
8-cell
Morula
Blastocyst
H3K27ac level
+TSA
b
Control
+TSA
c
***
Development (%)
Control
+TSA
d
p < 2.2e−16
Cell Number
Control
+TSA
g
Control
+TSA
TSS
TES
e
DNA
H3K27ac
merge
control
+TSA
0 µm 75
f
p < 2.2e−16
Relative intensity
Control
+TSA
h
Normalized RPKM
Control
+TSA
TSS
TES
i
TSA rep2
TSA rep1
Control rep2
Control rep1
j
upregulated
downregulated
−log10 (pvalue)
log2 FoldChange
Zfp982
Onecut2
Lratd1
Gm20765
Duxf3
Smarca2
Sox11
Sp9
Sox1
Eif4e1b
Gata2
Usp21
k
H3K27ac
Control
+TSA
RNA
Log2 FC (TSA/Con)
late 2-cell
minor ZGA genes
TSS
TES

**Fig. S6 Failure of zyH3K27ac removal resulted in excessive minor ZGA. a** Schematic presentation of the experimental TSA treatment protocol. **b** Representative images and (**c**) developmental rates of control and TSA-treated embryos. Data are from three independent experiments. ***$P < 0.001$ (Student's t-test). Scale bars, 50 μm. **d** Quantification of cells in control and TSA-treated embryos. Dots represent cell numbers in a single embryo. **e** Representative confocal images of control and TSA-treated embryos stained with H3K27ac antibody. Scale bars, 75 μm. **f** Quantification of H3K27ac fluorescence intensity in control and TSA-treated embryos. Each dot represents a single nucleus. **g** Heatmap showing zyH3K27ac signals ranked by their relative change after TSA treatment. **h** Metaplot of H3K27ac signals (Z-score normalized) in control and TSA-treated late 2-cell embryos. **i** Unsupervised clustering of gene expression among control and TSA-treated embryos at the late 2-cell stage. **j** RNA-seq analysis results for control and TSA-treated late 2-cell mouse embryos. Volcano plots show gene expression changes. Yellow and blue dots indicate significantly upregulated (fold change > 1) and downregulated (fold change < −1) genes ($P < 0.05$). **k** Heatmap (left) showing H3K27ac signals ranked by their relative changes after TSA treatment. Heatmap (right) shows that minor ZGA genes were upregulated between control and TSA-treated embryos. Mean values of two biological replicates were scaled and are represented as Z scores.

a

Gene expression during preimplantation development (FPKM)

| Genes | Zygote | early 2-cell | middle 2-cell | late 2-cell | 4-cell | 8-cell | Morula | Blastocyst |
|---|---|---|---|---|---|---|---|---|
| *Sirt1* | 13.8 | 19.8 | 185.3 | 169.9 | 116.9 | 46.5 | 36.4 | 21.6 |
| *Sirt2* | 4.8 | 5.0 | 4.6 | 5.2 | 15.4 | 18.5 | 16.1 | 26.5 |
| *Sirt3* | 10.5 | 9.7 | 7.9 | 3.6 | 6.3 | 5.5 | 1.2 | 14.9 |
| *Sirt4* | 11.1 | 7.6 | 13.3 | 7.1 | 5.2 | 0.4 | 2.3 | 2.2 |
| *Sirt5* | 20.7 | 23.1 | 10.9 | 5.7 | 8.5 | 17.2 | 12.6 | 4.7 |
| *Sirt6* | 43.7 | 27.2 | 6.6 | 28.7 | 79.8 | 65.8 | 90.9 | 44.0 |
| *Sirt7* | 29.6 | 9.2 | 28.2 | 61.3 | 69.7 | 26.0 | 18.6 | 11.0 |

b

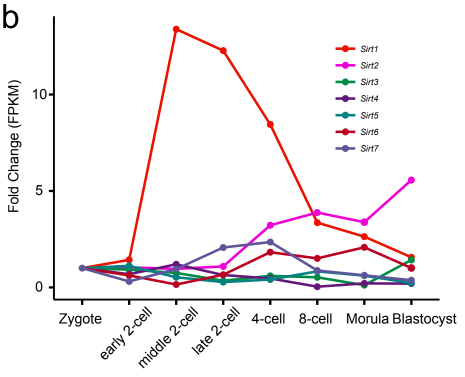


c

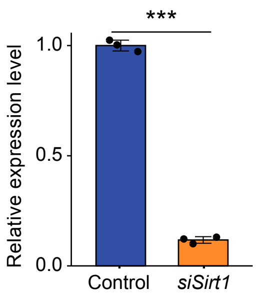


e

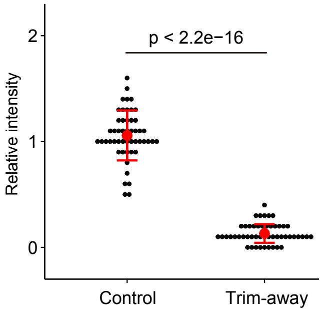


d

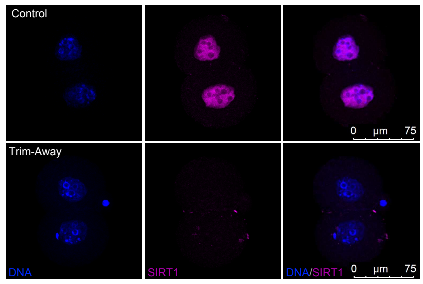


f

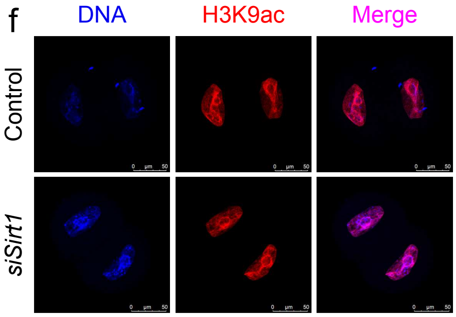


g

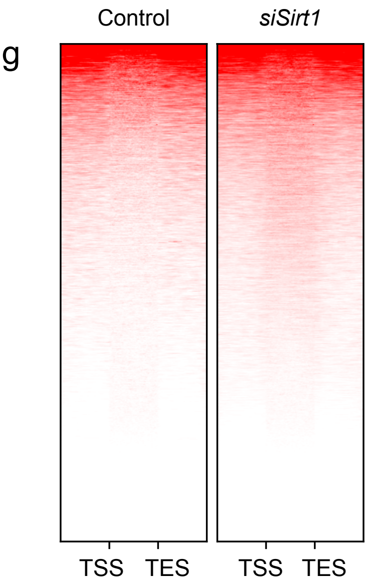


h

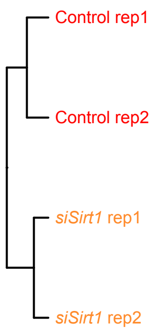


i

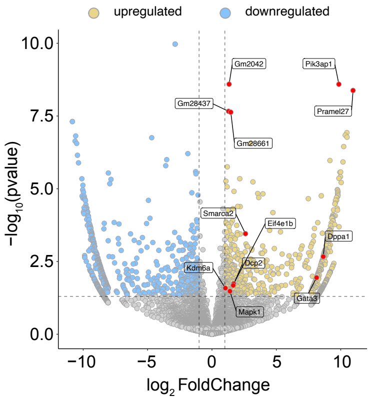


j

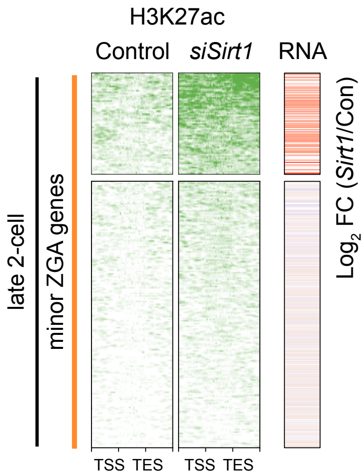


k

Domain architecture comparison of SIRT1 between human and mouse

| Score | Expect | Method | Identities | Positives | Gaps | Frame |
|---|---|---|---|---|---|---|
| 1148 bits(2970) | 0.0() | Compositional matrix adjust. | 632/748(84%) | 671/748(89%) | 12/748(1%) | |

```
Human 1     MADEAALALQPGGSPSAAGADREAASSPAGEPLRKRPRRDGPGLERSPGEPGGAAPEREV  60
            MADE ALALQ  GS  +A A  EAAS PA EPLRKRPRRDGPGL RSPGEP  A
Mouse 1     MADEVALALQAAGS-PSAAAAMEAASQPADEPLRKRPRRDGPGLGRSPGEPSAAVAPAAA  59

Human 61    PAAARGCPGAAAAALWREAEAEAAAAGGEQEAQATAAAGEGDNGPGLQGPSREPPLADNL  120
               A      AA AALWREA   AA+A  E+EA ATA AG+GDNG GL+   REP  AD+
Mouse 60    GCEAAS--AAAPAALWREAAGAAASA--EREAPATAVAGDGDNGSGLR---REPRAADDF  112

Human 121   YDEDDDDEGEEEEEAAAAAIGYRDNLLFGDEIITNGFHSCESDEEDRASHASSSDWTPRP  180
             D++ ++E E    AAAAAIGYRDNLL  D ++TNGFHSCESD++DR SHASSSDWTPRP
Mouse 113   DDDEGEEEDEAAAAAAAAAIGYRDNLLLTDGLLTNGFHSCESDDDDRTSHASSSDWTPRP  172

Human 181   RIGPYTFVQQHLMIGTDPRTILKDLLPETIPPPELDDMTLWQIVINILSEPPKRKKRKDI  240
            RIGPYTFVQQHLMIGTDPRTILKDLLPETIPPPELDDMTLWQIVINILSEPPKRKKRKDI
Mouse 173   RIGPYTFVQQHLMIGTDPRTILKDLLPETIPPPELDDMTLWQIVINILSEPPKRKKRKDI  232

Human 241   NTIEDAVKLLQECKKIIVLTGAGVSVSCGIPDFRSRDGIYARLAVDFPDLPDPQAMFDIE  300
            NTIEDAVKLLQECKKIIVLTGAGVSVSCGIPDFRSRDGIYARLAVDFPDLPDPQAMFDIE
Mouse 233   NTIEDAVKLLQECKKIIVLTGAGVSVSCGIPDFRSRDGIYARLAVDFPDLPDPQAMFDIE  292

Human 301   YFRKDPRPFFKFAKEIYPGQFQPSLCHKFIALSDKEGKLLRNYTQNIDTLEQVAGIQRII  360
            YFRKDPRPFFKFAKEIYPGQFQPSLCHKFIALSDKEGKLLRNYTQNIDTLEQVAGIQRI+
Mouse 293   YFRKDPRPFFKFAKEIYPGQFQPSLCHKFIALSDKEGKLLRNYTQNIDTLEQVAGIQRIL  352

Human 361   QCHGSFATASCLICKYKVDCEAVRGDIFNQVVPRCPRCPADEPLAIMKPEIVFFGENLPE  420
            QCHGSFATASCLICKYKVDCEAVRGDIFNQVVPRCPRCPADEPLAIMKPEIVFFGENLPE
Mouse 353   QCHGSFATASCLICKYKVDCEAVRGDIFNQVVPRCPRCPADEPLAIMKPEIVFFGENLPE  412

Human 421   QFHRAMKYDKDEVDLLIVIGSSLKVRPVALIPSSIPHEVPQILINREPLPHLHFDVELLG  480
            QFHRAMKYDKDEVDLLIVIGSSLKVRPVALIPSSIPHEVPQILINREPLPHLHFDVELLG
Mouse 413   QFHRAMKYDKDEVDLLIVIGSSLKVRPVALIPSSIPHEVPQILINREPLPHLHFDVELLG  472

Human 481   DCDVIINELCHRLGGEYAKLCCNPVKLSEITEKPPRTQKELAYLSELPPTPLHVSEDSSS  540
            DCDVIINELCHRLGGEYAKLCCNPVKLSEITEKPPR QKEL +LSELPPTPLH+SEDSSS
Mouse 473   DCDVIINELCHRLGGEYAKLCCNPVKLSEITEKPPRPQKELVHLSELPPTPLHISEDSSS  532

Human 541   PERTSPPDSSVIVTLLDQAAKSN-DDLDVSESKGCMEEKPQEVQTSRNVESIAEQMENPD  599
            PERT P DSSVI TL+DQA  +N +DL+VSES  C+EEKPQEVQTSRNVE+I   +ENPD
Mouse 533   PERTVPQDSSVIATLVDQATNNNVNDLEVSES-SCVEEKPQEVQTSRNVENI--NVENPD  589

Human 600   LKNVGSSTGEKNERTSVAGTVRKCWPNRVAKEQISRRLDGNQYLFLPPNRYIFHGAEVYS  659
             K VGSST +KNERTSVA TVRKCWPNR+AKEQIS+RL+GNQYLF+PPNRYIFHGAEVYS
Mouse 590   FKAVGSSTADKNERTSVAETVRKCWPNRLAKEQISKRLEGNQYLFVPPNRYIFHGAEVYS  649

Human 660   DSEDDVLSSSSCGSNSDSGTCQSPSLEEPMEDESEIEEFYNGLEDEPDVPERAGGAGFGT  719
            DSEDDVLSSSSCGSNSDSGTCQSPSLEEP+EDESEIEEFYNGLED+ + PE AGG+GFG
Mouse 650   DSEDDVLSSSSCGSNSDSGTCQSPSLEEPLEDESEIEEFYNGLEDDTERPECAGGSGFGA  709

Human 720   DGDDQEAINEAISVKQEVTDMNYPSNKS  747
            DG DQE +NEAI+ +QE+TD+NYPS+KS
Mouse 710   DGGDQEVVNEAIATRQELTDVNYPSDKS  737
```

**Figure S7. Effect of *Sirt1* knockdown on zyH3K27ac removal. a, b** Sirtuin family expression patterns according to RNA-seq data in early mouse embryos. Results of (**c**) qPCR and (**d**) immunofluorescence analysis of late 2-cell *Sirt1* knockdown embryos, validating the *Sirt1* knockdown efficiency results. **e** Quantification of SIRT1 the fluorescence intensity of control and *Sirt1* knockdown in late 2-cell embryos. Each dot represents a single nucleus. **f** Representative confocal images of control and *Sirt1*-knockdown embryos stained with H3K9ac antibody. Scale bars, 50 μm. **g** Heatmap showing zyH3K27ac signals ranked by their relative change after *Sirt1* knockdown in late 2-cell embryos. **h** Unsupervised clustering of gene expression among control and *Sirt1* knockdown embryos at the late 2-cell stage. **i** RNA-seq analysis results for control and *Sirt1* knockdown in late 2-cell mouse embryos. Volcano plots show gene expression changes. Yellow and blue dots indicate significantly upregulated (fold change > 1) and downregulated (fold change < −1) genes ($P < 0.05$). **j** Heatmap (left) showing H3K27ac signals ranked by their relative change after *Sirt1* knockdown. Heatmap (right) shows that minor ZGA genes were upregulated between control and *Sirt1*-knockdown embryos. Mean values of two biological replicates were scaled and are represented as Z scores. **k** SIRT1 proteins are conserved between human and mouse, yellow boxes indicate functional domains.
